# Respiratory Complex I Regulates Dendritic Cell Maturation in Explant Model of Human Tumor Immune Microenvironment

**DOI:** 10.1101/2023.05.10.539944

**Authors:** Rita Turpin, Ruixian Liu, Pauliina M. Munne, Aino Peura, Jenna H. Rannikko, Gino Philips, Bram Boeckx, Natasha Salmelin, Elina Hurskainen, Ilida Suleymanova, Elisa M. Vuorinen, Laura Lehtinen, Minna Mutka, Panu E. Kovanen, Laura Niinikoski, Tuomo Meretoja, Johanna Mattson, Satu Mustjoki, Päivi Saavalainen, Andrei Goga, Diether Lambrechts, Jeroen Pouwels, Maija Hollmén, Juha Klefström

## Abstract

Combining cytotoxic chemotherapy or novel anticancer drugs with T-cell modulators holds great promise in treating advanced cancers. However, the response varies depending on the tumor immune microenvironment (TIME). Therefore, there is a clear need for pharmacologically tractable models of the TIME to dissect its influence on mono- and combination treatment response at the individual level. Here we establish a Patient-Derived Explant Culture (PDEC) model of breast cancer, which retains the immune contexture of the primary tumor, recapitulating cytokine profiles and CD8+ T cell cytotoxic activity. We explored the immunomodulatory action of a synthetic lethal BCL2 inhibitor venetoclax + metformin drug combination *ex vivo*, discovering metformin cannot overcome the lymphocyte-depleting action of venetoclax. Instead, metformin promotes dendritic cell maturation through inhibition of mitochondrial complex I, increasing their capacity to co-stimulate CD4+ T cells and thus facilitating anti-tumor immunity. Our results establish PDECs as a feasible model to identify immunomodulatory functions of anticancer drugs in the context of patient-specific TIME.

## INTRODUCTION

While a common denominator of cancer is a dysregulation of the tumor immune microenvironment (TIME), the actual composition and function of the TIME is both tumor type- and patient-specific^1^. Characterizing the TIME is improving the stratification of patients who may respond to immunomodulatory monotherapies, like anti-CTLA-4 or anti-PD-1/PD-L1^2^. However, it is increasingly clear that the TIME also has a strong role in influencing the outcome of cytotoxic chemo or targeted therapies which were not meant to, or do not affect immune cells directly. While the release of tumor antigens by tumor toxic compounds could trigger a beneficial immune response^3,4^, the drug toxicity also targets mitotically active immune cells with potentially negative effects on anti-tumor immune response^5, 6^. Furthermore, chemo-induced immunogenic cell death can, in certain circumstances, act as a trigger for the recruitment of pro-tumor macrophages^7^, which can reduce the sensitivity of cancer cells to paclitaxel, etoposide, and doxorubicin^8^.

Recent clinical studies in breast cancer (BC) have also shown that targeting the immune system through the PD-1/PD-L1 pathway alone is not effective^9–11^, whereas a combination of paclitaxel + anti-PD-L1, designed to target the tumor and the TIME simultaneously, has provided clinical evidence of efficiency to support approval ^12, 13^. In fact, the concept of targeting tumor cells and immune cells simultaneously is so widely tested as a treatment modality for different cancer types that nearly 90% of current PD-1/PDL1-targeted trials include a combination therapy^14^.

These notions highlight that defining anticancer drug effects on not only tumor cells, but also on the TIME is crucial to understand how the drug’s action translates into efficacy in the context of a heterogeneous tumor microenvironment. A better understanding of which immune cell types are activated, depleted, or otherwise impacted under the treatment, would provide important clinical trajectories especially for a choice of right combination immunotherapies in a personalized treatment setting ^15, 16^.Synthetic lethality (SL) is a concept that describes the selective killing of cancer cells which harbor specific alterations of an oncogenic or tumor suppressor pathway, with a drug that is toxic to cancer cells due to specific drug-sensitizing mutations, but is much less toxic to normal cells that are lacking the mutations ^17^. The MYC gene is amplified, or the MYC-encoded protein is elevated through other mechanisms, in up to 70% of human cancers^18^. The elevated or deregulated MYC levels drive many oncogenic processes, including metabolic reprogramming and non-stop cell cycle progression, but MYC also sensitizes cells to diverse inducers of extrinsic or intrinsic programmed cell death pathways^19–22^. These early findings have laid the conceptual foundation for MYC-dependent synthetic lethal (MYC SL) therapeutic strategies, which seek to specifically harness MYC generated vulnerability pathways as opposed to drugging the MYC protein directly^23–25^. A previous study exploring therapeutic opportunities through the MYC SL concept revealed that in several mouse models of MYChigh breast cancer, the MYChigh tumors *in vivo* are specifically vulnerable to a combination treatment with venetoclax and metformin. Venetoclax is a BH3-mimetic that blocks the anti-apoptotic B cell lymphoma-2 (BCL-2) protein^26^ and metformin is a commonly prescribed drug for type 2 diabetes^27^. In particular, the treatment of syngrafted Wap-Myc tumors in mice with the venetoclax+metformin (VeM) combination results in cessation of tumor growth and the addition of anti-PD-1 checkpoint inhibitor to the treatment regimen results in a persistent treatment response - with no tumors growing in the mice even after drug withdrawal. While these observations are consistent with the idea of immunogenic cell death as a mechanism of VeM *in vivo*, emerging evidence also suggests an important immunomodulatory function for metformin; it can maintain high cytotoxic T lymphocyte (CTL) activity in tumor cells^28^ through enhancing the antiapoptotic abilities of CD8+ T cells and downmodulating PD-1/PD-L1^29, 30^. Therefore, the anticancer effects observed with the VeM combination raises the interesting question whether metformin only acts by boosting cancer cell death, or whether it somehow modulates an immune response directly.

We previously developed a method to grow 3D cultures of intact fragments of primary human patient-derived breast and breast cancer tissue. The Patient-Derived Explant Culture (PDEC) model offers many advantages over conventional reductionist and artifact-prone cell co-culture and rodent models, and it has provided new insights into the biology of breast cancer subtypes, as well as mechanisms of treatment regimens in the context of authentic human breast tumor tissue^27, 31–33^. Since PDECs come directly from surgery, they contain viable immune cells. We considered that PDECs could offer a unique method to simultaneously investigate the effects of VeM on tumor cells and tumor resident immune cells in *ex vivo* conditions. Here, we first show that PDECs maintain the immune composition and baseline immune activity of the primary breast tumors. In addition, we demonstrate that the cytolytic activity of PDEC containing T cells can be activated with a direct T cell activator, anti-CD3/CD28/CD2, but not via the PD-1/PDL1 mechanism. Venetoclax depleted tumor-infiltrating lymphocytes in PDECs, as previously observed in mice^34^, an effect that was not counteracted by metformin. However, we report that metformin surprisingly promotes human dendritic cell (DC) maturation by altering the immune cell metabolism through the inhibition of respiratory complex I (CI). The metformin-induced DC maturation can trigger CD4+ T cell proliferation, suggesting the inhibition of CI or other sites of mitochondrial respiration as a potential new therapeutic strategy to enhance immunotherapy.

## RESULTS

### PDECs maintain the immune contexture and baseline immune activity of the primary breast tumor

To define whether PDECs preserve components of the primary TIME, we investigated the presence of CD45+ leukocytes in the tumor explant cultures using confocal immunofluorescence microscopy. The analysis of explants from five different patients after three days of ex vivo culture showed the presence of CD45+ leukocytes in all studied cultures (Fig 1a). To determine how well PDECs recapitulate the immune cell composition of the primary tumor sample, a workflow was designed to compare the general composition of tumor-infiltrating leukocytes (TILs) of primary tumors, with their corresponding PDECs up to one week in culture (Fig 1b). First, the immune cell composition of two donors was analyzed using single cell RNA sequencing of sorted CD45+ leukocytes comparing primary tumor material to explants grown for one week. The results demonstrated that myeloid cells, B cells, NK (Natural Killer) cells, CD8+ T cells, and CD4+ T cells were preserved in PDECs in similar proportions as the primary tumor, with no significant differences between cell types (Fig 1c-e; **Supplementary Fig 1a-b**). Second, while Nanostring nSolver Gene expression profiling at three days in culture revealed an overall decrease in tumor-infiltrating leukocytes within explants (Fig 1f), the composition of the TILs which includes B cells, cytotoxic cells, macrophages, neutrophils, NK cells, T cells (including Th1, Treg, and CD8+) was surprisingly similar (Fig 1g). We did, however, observe a decrease in the proportion of mast cells, and an increase in the proportion of dendritic cells (DCs) (Fig 1g). Third, flow cytometric comparison of CD45+ leukocytes of the primary tumors and PDECs cultured ex vivo for one week confirmed that the relative numbers of lymphocytes (CD4+ and CD8+ T cells, NK, NKT) were not significantly altered (Fig 1h).

**Figure. 1.**
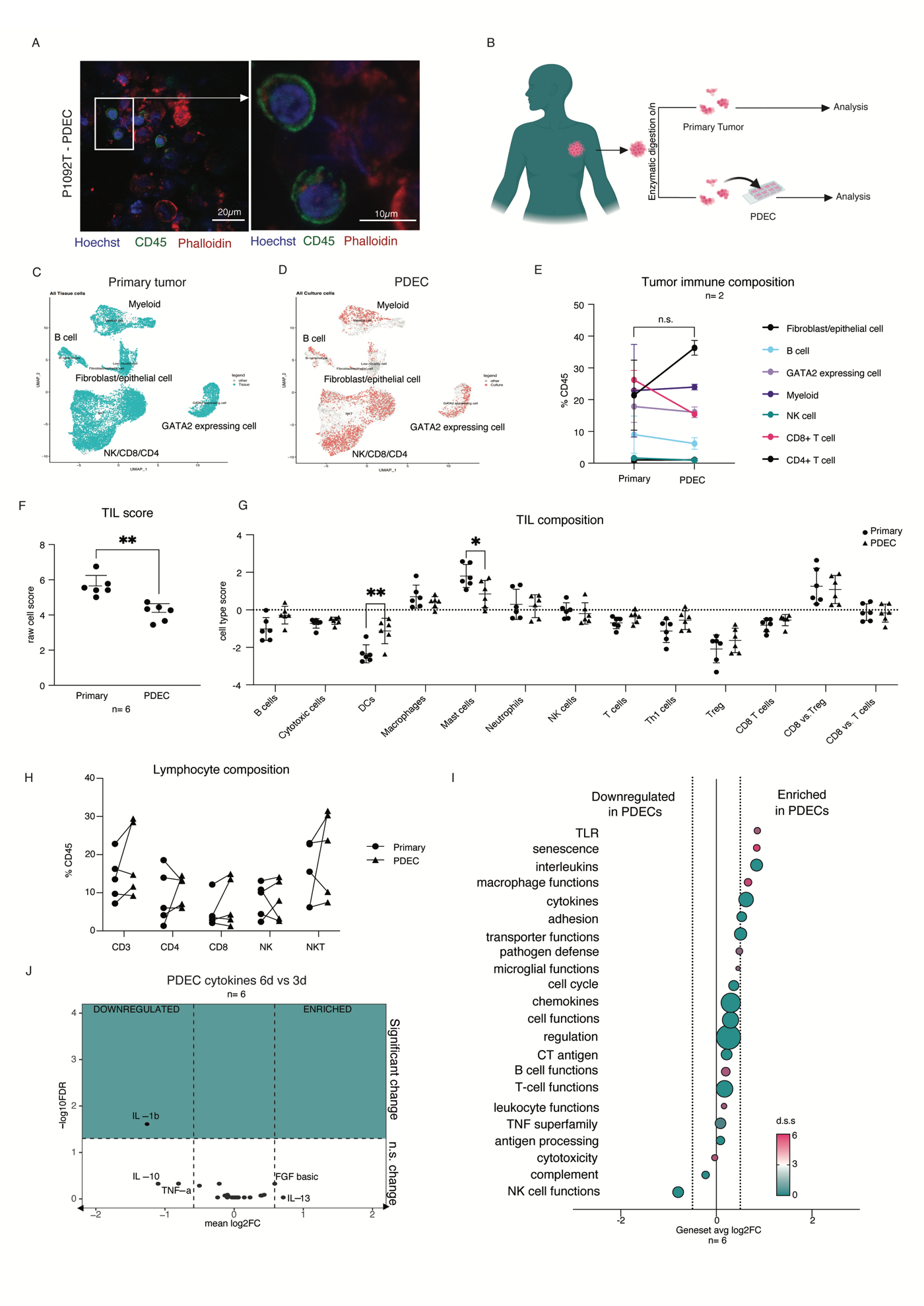
PDECs maintain the immune contexture and baseline immune activity of primary breast tumor. **a,** immunofluorescent staining of CD45 and F-actin in PDECs after 72hrs in culture. P1092T refers to a tumor sample from patient 1092. **b,** Schematic representation of workflow comparing primary tumor tissue to cultured tumor tissue (PDEC). **c,** Single-cell RNA sequencing UMAPs after data integration from n=2 primary tumors, **d**, and the corresponding PDECs. **e,** immune cell composition of primary tumors compared with the two corresponding PDECs. No significant changes detected between any immune subtypes using the MASC algorithm **f**, gene expression profiling of leukocytes of biologically independent primary tumors and their corresponding PDECs shows a decrease in TIL gene expression (p= 0.0013) after 72hrs in culture (n=6). Statistics were done with paired t test with two-tailed p value. **g,** Immune cell composition obtained from gene expression profiling of samples from (f) normalized to nSolver tumor infiltrating leukocyte gene signature. Dendritic cell (p=0.0032) and mast cell (p=0.0481) numbers were significantly affected in culture. Statistics are 2way ANOVA with Sidaks multiple comparison test. **h,** flow cytometry analysis of CD3+ T cells, CD4+ T helper cells, CD8+ T effector cells, NK, and NKT cells from primary tumors and corresponding PDECs (n=8) with no significant differences between primary tumor and PDECs **i,** estimation of immune cell activity pathways from nanostring gene expression profiling normalized to TIL numbers. Pink fuchsia color indicates directional significance (t-statistic for each gene against each covariate) Pathways considered significantly different are those with an average log2fold change of > 0.5 in addition to statistically significant overexpression as determined by the directional significance score above 0. **j**, Cytokine profiling of explant media at 72hrs, and 144hrs. Cytokines below −0.58 log2FC are downregulated more than 1.5-fold, and those with -log10FDR of 1.3 or greater are significant. Cytokine statistics were computed with one-sample t-test for deviance from 0. p-values were adjusted by FDR method. All data are presented as mean values +/− SD.

To determine the baseline activity status of primary tumor vs. PDEC TILs, the gene expression profiling samples were processed to remove systematic differences in TIL numbers, and further analyzed for cell and immune activity genesets (Fig 1i). The basal activity of pathways including NK cell functions, T cell functions, B cell functions, cytokines and interleukins was similar to the primary tumor sample **(Supplementary Fig 1c-d**). Only a few pathways, including TLR, senescence, and macrophage functions, were significantly different but we note that their corresponding profiles consisted of a small number of genes **(supplementary Fig 1d).** Additionally, longitudinal cytokine profiling up to one week revealed a significant decrease in only one cytokine, 1L-1b, while the other 26 were not significantly changed (Fig 1j). These results suggest that the PDEC cultures themselves have little effect on the baseline activity of the immune cells. Overall, these results indicate that the PDEC model preserves the TIME of the primary patient tumor during a seven-day culture period.

### Resident tumor-infiltrating T cells can be activated to kill tumor cells ex vivo

The presence of CD45+ cells, including cytolytic cells of both innate (NK, NKT) and acquired immunity (CD8+ T cells), led us to ask whether these cells could be functionally activated. We used a commercially available soluble antibody complex, Anti-CD3/CD28/CD2, which cross-links the surface ligands CD2, CD3, and CD28 on T cells to provide stimulatory and co-stimulatory signals needed for robust T cell activation, herein referred to as anti-CD3/CD28/CD2. We compared anti-CD3/CD28/CD2 to therapeutically relevant antibodies for programmed cell death 1 (PD-1) and its ligand (PD-L1) (programmed death ligand 1) to detect the “maximum” T cell response in PDECs, and to determine whether in our PDEC model the PD-1-PD-L1-axis is a critical signaling pathway that limits the physiological activity of resident T cells^35^. We treated the explants for 72hrs and measured general cell death with the CellTox green assay, observing a statistically significant increase in cell death upon anti-CD3/CD28/CD2 treatment whereas no cell death was observed following treatment with anti-PDL1 (Fig 2a, b). We randomly chose three primary tumor samples from the CellTox experiment and performed multiplex immunohistochemistry (IHC) to confirm that the samples contained immune cells, and that the cell death was a result of T cell cytotoxicity (Fig 2c; **Supplementary Fig 2a-b**). To further verify that the anti-CD3/CD28/CD2-mediated cell death signal potentially came from tumor cells, we performed flow cytometric analysis of PDECs, and quantified absolute numbers of CD45- non-hematopoietic cell populations, which were mostly tumor cells, but also stromal cells like fibroblasts. Here we also observed a reduction of tumor cells after T cell activation (Fig 2d).

**Figure. 2.**
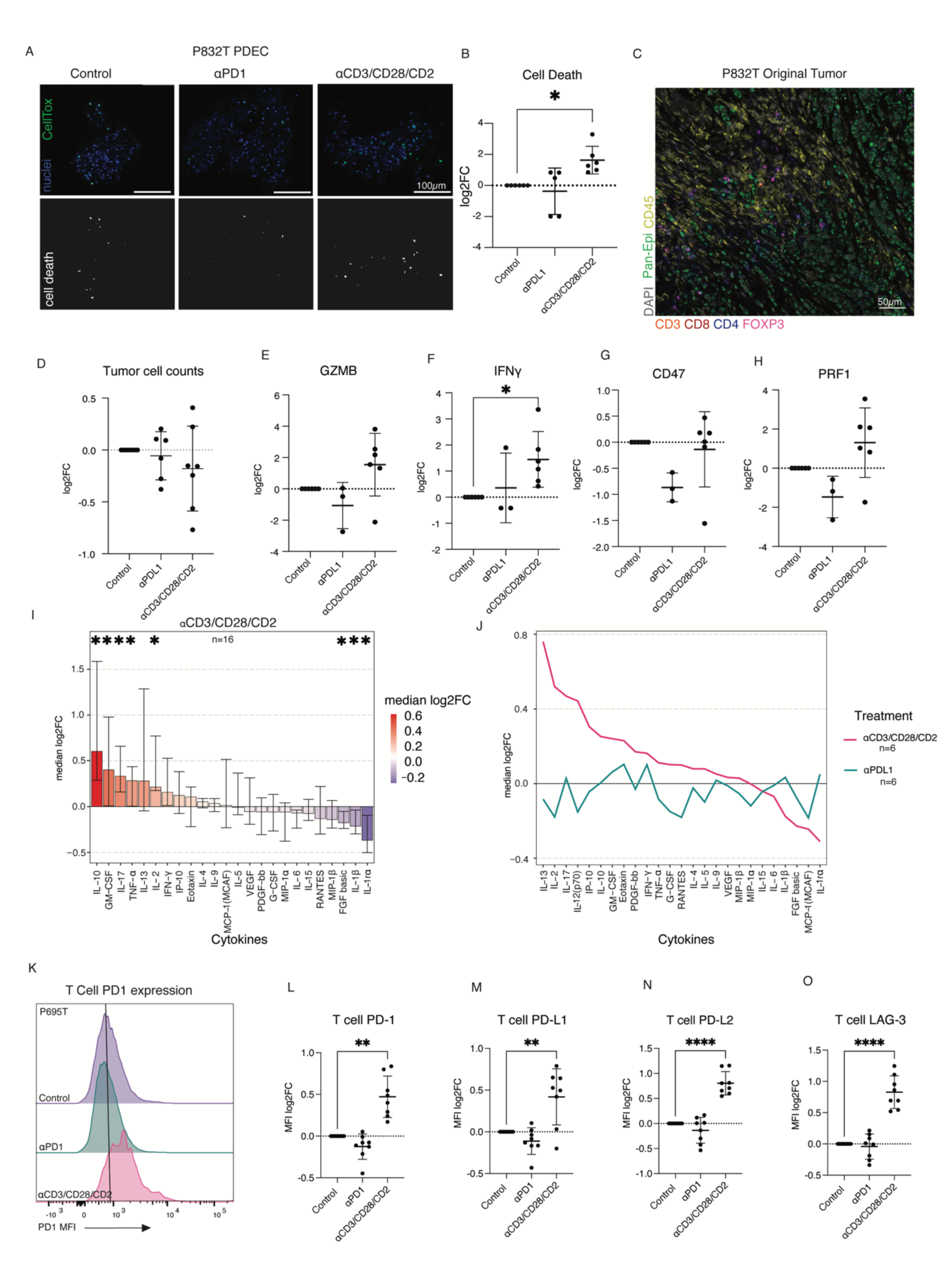
Resident tumor-infiltrating T cells can be activated to kill tumor cells ex vivo. **a,** CellTox staining of P832T PDEC following treatment with a-PD-1 and anti-CD3/CD28/CD2. **b,** Quantification of cell death from IF images corresponding to n=6 biologically independent PDECs, showing significant cell death following anti-CD3/CD28/CD2 treatment (p=0.0321) quantified with a one-way ANOVA with Fishers exact test. **c,** immune infiltration of primary tumor from which PDECs in Fig 3a. were derived from. **d,** Tumor cell (CD45-) counts in n=7 biologically independent PDECs (n=6 for aPDL1) following aPDL1 and antiCD3/CD28/CD2 treatment as the log2FC of the absolute cell numbers normalized the control **e-h,** qPCR analysis of *GZMB, IFNɣ* (p= 0.0208)*, CD47, PRF1* relative to the control. n= 3 for aPDL1, n=6 for control and anti-CD3/CD28/CD2-treated. Statistical significance was tested with a one-way ANOVA with Fishers exact test. Data are presented as mean values +/− SD. **i,** multiplex cytokine profiling of PDECs after 72hrs anti-CD3/CD28/CD2 treatment. n=16. Each cytokine is tested with a one-sample non-parametric wilcox.test for deviance from 0 (no change) and p-values adjusted by FDR method. Cytokines ordered by median log2FC and plotted with error bars ranging the interquartile range. **j,** Comparison of median log2FC of cytokines between anti-CD3/CD28/CD2 and aPDL1 treatments in PDECs. n=6. **k,** flow cytometry representation of median fluorescence intensity of checkpoint marker, PD-1, on CD3+ T cells in control, aPD1 and anti-CD3/CD28/CD2-treated PDEC of P695T. **l-o,** graphs quantifying median surface expression of T cell checkpoint proteins relative to control. Statistical significance was tested with significant increase in PD-1 (p=0.0010), PD-L1 (p=0.0096), PD-L2 (p=<0.0001), and LAG-3 (p=<0.0001) in response to anti-CD3/CD28/CD2. Statistical significance was tested with a one-way ANOVA with Fishers exact test. Data are presented as mean values +/− SD.

Consistent with an increase in cytotoxic activity leading to cell death, qPCR analysis of PDECs following anti-CD3/CD28/CD2 treatment revealed a statistically significant increase in IFNɣ expression, and a trend for elevated mRNA expression of GZMB and PRF1 (Fig 2e-h). Both PRF1 and GZMB are expressed by cytotoxic T cells and NK cells upon activation, while IFNɣ is a signature proinflammatory cytokine typically used to measure immune activation^36, 37^. The cell death and mRNA results in response to anti-CD3/CD28/CD2, along with the fact that anti-PDL1 monotherapy has no positive effect on tumor cell death, or the transcription of *IFNɣ, GZMB, or PRF1*, reinforce the idea that the cell death seen previously is likely a result of TCR-triggering, and the downstream effects of T cell activation.

We profiled cytokines of T cell activation and inflammation from PDEC supernatant after 72hrs of anti-CD3/CD28/CD2 (n=16), and anti-PDL1 (n=6) treatments. The results highlight the ability of PDECs to reflect patient heterogeneity, while still capturing the general trend of cytokines expected after robust T cell activation (Fig 2i-j). For example, we observed an increase in IL-2 which is secreted by CD4+ T cells upon antigen stimulation, and in IL-10, which is secreted in response to immune activation by several different cell types^38, 39^ (Fig 2i). In accordance with the previous qPCR findings, we observed that unlike anti-CD3/CD28/CD2 treatment, anti-PDL1 monotherapy did not deviate far from the control, suggesting negligible effect from the drug in terms of induced cytokine expression (Fig 2j). For T cell-specific analysis, CD3+ T cells were profiled directly for signs of exhaustion following treatment by flow cytometry, measuring the median fluorescent intensity of checkpoint ligands/receptors PD-L2, PD-L1, LAG3, and PD1 on the surface of the cells. Once again, anti-CD3/CD28/CD2, but not anti-PD-1, induced expression of these checkpoint molecules, suggestive of T cell activation (Fig 2k-o; **Supplementary Fig 2c**). The transcriptional and cytokine data support the general detection of immune activation within PDEC cultures, while IF and flow cytometry analysis of PDECs suggests that T cells are functionally activatable to kill tumor cells ex vivo.

Together, the presence of the primary tumor TIL repertoire within PDECs and the evidence for activatable cytolytic properties in the tumor CD8+ T cells demonstrate PDECs as a versatile model to study various aspects of tumor-immune tissue interactions and anticancer drug responses in context of the TIME.

### Assessment of combination therapy with venetoclax reveals lymphodepletion of T cells

Since the PDECs preserved the TIME, we recognized an opportunity to further explore and validate the specific responses of tumor cells and TIME to the venetoclax+metformin (VeM) combination, which shows strong in vivo antitumor activity in several mouse models of MYC-driven aggressive breast cancer^27^. Clinically, venetoclax has been observed as a lymphodepleting agent as it induces apoptosis in BCL-2 reliant subsets of T cells^34^. The lymphodepleting activity is a potential clinical concern, as it could negatively impact immune system functions and reduce the opportunities to combine venetoclax with immune-modulating therapies. We first tested the idea that metformin component of VeM perhaps affords protection to immune cells against the lymphodepleting actions of venetoclax, which could explain the better in vivo effect of VeM in comparison to single treatments^27^. In parallel with VeM, we treated PDECs with clinically relevant paclitaxel, a chemotherapy used to treat primary breast tumors, often in combination with other drugs (Fig 3a).

**Figure. 3.**
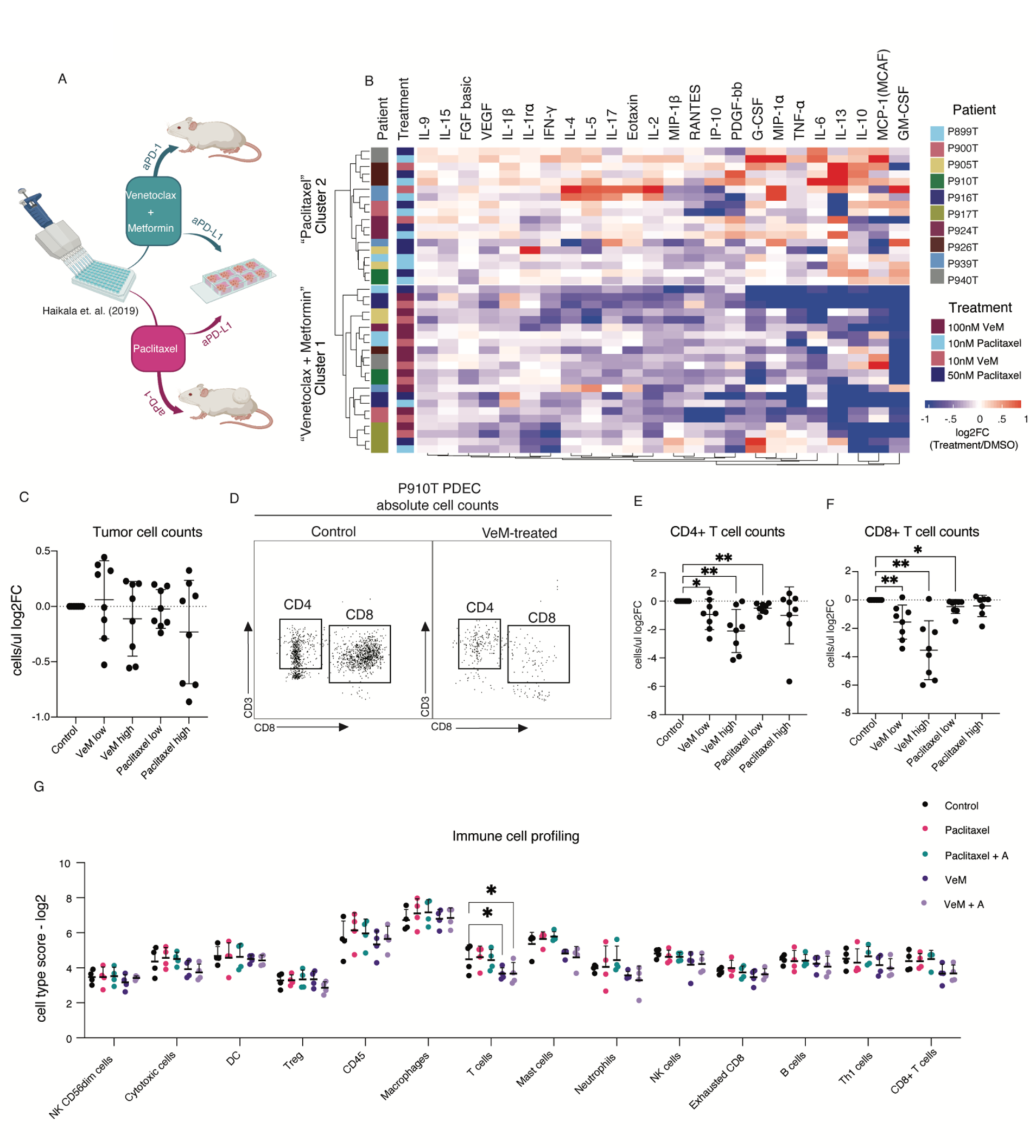
Assessment of combination therapy with venetoclax reveals lymphodepletion of T cells. **a,** schematic summary of previous finding^27^ in WAP-Myc mouse model of MYC-driven breast cancer where venetoclax+metformin+aPD1 treatment in mice resulted in durable antitumor immunity. Here, the VeM treatment along with paclitaxel were profiled in PDECs. **b,** Unsupervised hierarchical clustering (Euclidean distance, complete linkage) of PDECs based on cytokine secretion following venetoclax + metformin and paclitaxel treatments (n = 10 biologically independent PDECs). **c,** CD45-tumor cell viability after VeM or paclitaxel treatments **d,** representative image of quantifying cell viability through flow cytometry for panels e-f **e,** Low (10nM venetoclax + 10mM metformin) (p=0.0428) and high (100nM ventoclax + 10mM metfomin) (p=0.0059) concentrations of VeM negatively impact CD4 T cell viability (CD45+ CD3+ CD56-CD4+) **f,** Low (p=0.0082) and high (p=0.0019) concentrations of VeM negatively impact T effector (CD45+, CD3+, CD56-, CD8+) viability **g,** nanostring gene expression profiling of projected cell type scores following treatment of n=4 biologically independent PDECs. High VeM (100nM ventoclax + 10mM metfomin) p=0.0434) without and with aPD-L1 (p=0.0449) negatively impact the T cell score. Statistical significance was tested with a two-way ANOVA with Fishers LSD. All data are presented as mean values +/− SD.

Unsupervised clustering of cytokines of 10 patient samples revealed tight clustering by treatment. We observed a consistent downregulation of most T-cell-associated cytokines (as determined in Fig 2i) in response to VeM, irrespective of MYC-status, suggestive of a depletion of lymphocytes (Fig 3b, **supplementary Fig 3a**). Flow cytometry analysis of CD45-cells, mainly tumor cells, showed some variation in the level of apoptotic responses to VeM between independent PDEC samples (Fig 3c,d), whereas every VeM treated sample showed a noticeable drop in the number of CD4+ and CD8+ T cells, with a more significant decrease in the numbers of CD8+ T cells (Fig 3e,f). Although we failed to see tumor cytotoxicity in PDECs after paclitaxel treatment, likely due to a delay between the drug’s primary and secondary modes of action ex vivo^40^, we did notice an increase in inflammatory cytokines suggestive of inflammation (IL2, IL10, IL4, IL15) from a subset of patients. We took a transcriptional approach to broaden the variety of immune cell types we could observe simultaneously (Fig 3g). As with the cytokine profiling and flow cytometry we noticed a clear diminishing impact of VeM on T cells. These results led us to refute the initial working hypothesis suggesting that metformin protects BCL-2-reliant CTLs against the lymphodepleting effects of venetoclax since none of our experiments supported this notion.

### Metformin contributes to dendritic cell activation ex vivo and in vivo

The fact that metformin did not protect T cells from apoptosis following venetoclax treatment led us to hypothesize that the immunogenic benefits of metformin might derive from the immune cells that were spared within the PDEC tumor microenvironment; for example, dendritic cells, macrophages, and NK cells remained unchanged (Fig 3g). We looked into the gene expression profiles of VeM-treated explants from four patients, and found that many of the overexpressed genes, including *HMGB1, CD97, MAF, and LAMP-3,* could be attributed to the activation of antigen presenting cells (Fig 4a). A similar profile was observed for VeMA-treated patients, but not either paclitaxel or paclitaxel+ aPDL1 **(Supplementary Fig 4a-c**). A metascape analysis of the most significantly affected biological pathways following VeM treatment based on the overexpressed genes included: cytokine signaling in immune system, adaptive immune response, and regulation of mononuclear cell proliferation - suggesting that even with the depletion of T cells, there were still elements important for T cell response being upregulated (Fig 4b). These observations combined with an increase of macrophage-inflammatory protein 1-beta, and IL-17 which are secreted by innate lymphoid cells and monocyte-derived cells (Fig 4c), and the elevated expression of LAMP-3 on a larger set of patient samples **(Supplementary Fig 4d),** suggested we focus our investigation on the antigen presenting cells.

**Fig. 4.**
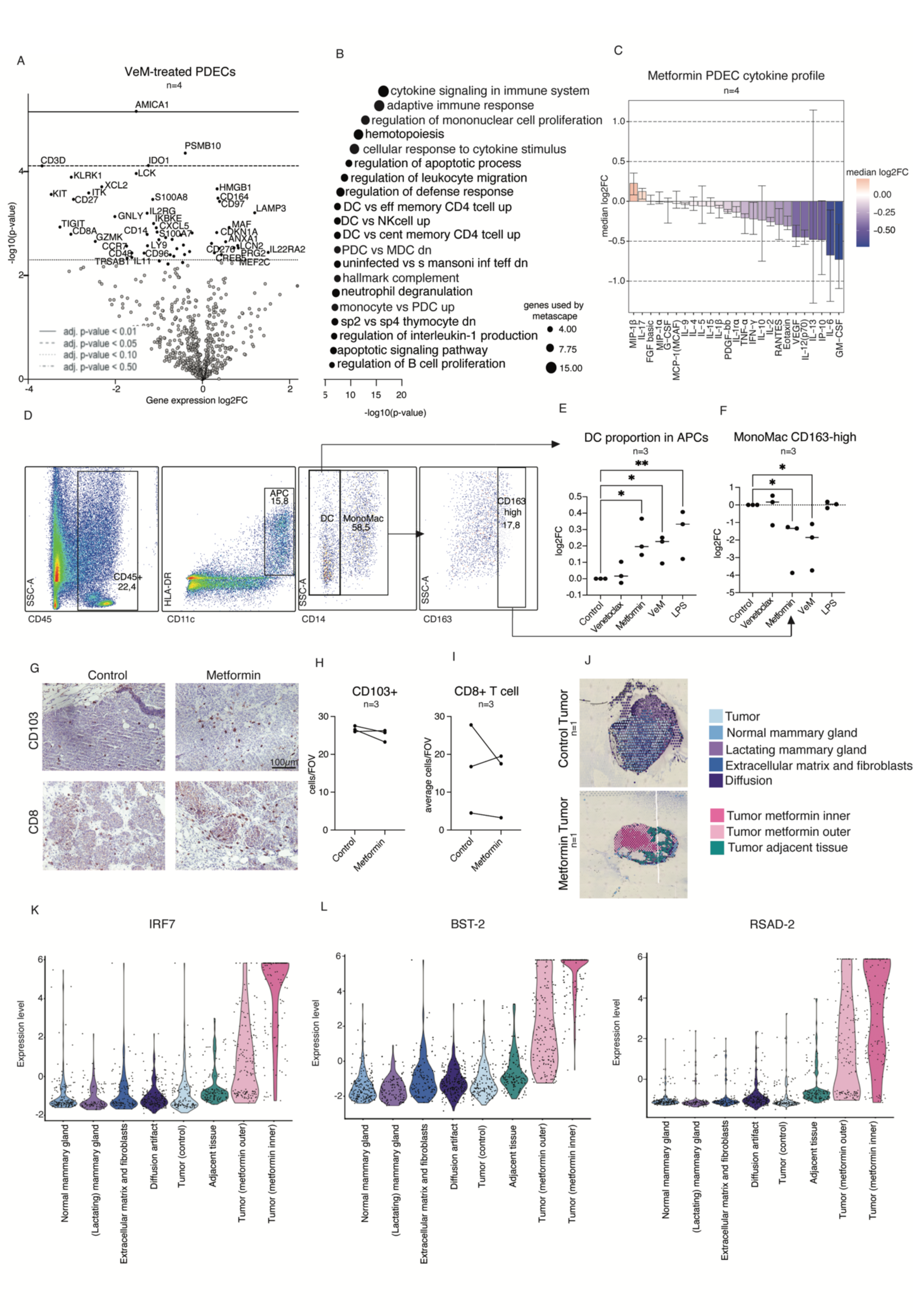
Metformin contributes to dendritic cell activation ex vivo and in vivo. **a,** Volcano plot of differentially expressed genes of n=4 VeM-treated PDECs. Adjusted p-value key described on the plot. **b,** metascape analysis of differentially expressed genes from (a) which are at least 20% upregulated and significant with a p-value of less than 0.05. The graph depicts terms which were significantly enriched after treatment. Ball size corresponds with the number of genes that overlapped with the gene signatures tested on metascape. **c,** cytokine profiling of PDECs following 10mM metformin treatment. Each cytokine was tested by one-sample non-parametric wilcox test for deviance from 0 (no change) and p-values adjusted by FDR method. n=4. **d,** flow cytometry gating strategy showing how dendritic cells, mono-macs, and high-CD163-expressing monomacs were determined. **e,** quantification of the proportion of DCs within the APCs **f,** quantification of the proportion of mono-macs expressing high levels of CD163 **g,** representative immunohistochemistry staining of CD103 dendritic cell activation marker and CD8 cytotoxic T cell marker in control and metformin-treated WapMyc mouse tumors (n=3 control, and n=3 metformin-treated mouse tumors). **h-i,** quantification of g. **j,** Control and metformin-treated mouse tumors from WapMyc mice with spatial annotations of tissue areas. Each circle is 10uM in diameter. **k-m,** mouse DC activation markers, *Bst2, Irf7*, and *Rsad2* in varying tumor regions as shown in j. All data are presented as mean values +/− SD. Statistical significance was tested with a one-way ANOVA with Fishers exact test.

In agreement with our gene expression profiling (Fig 3g.), we also did not detect a change in the number of antigen-presenting cells (APCs) following venetoclax, metformin, or VeM treatment with flow cytometry **(Supplementary Fig 4e).** However, with established surface marker panels for APCs^41, 42^, we found that metformin clearly increased the proportion of APCs with a dendritic cell (DC) -phenotype (CD11c+, HLA-DR+, CD14-) (Fig 4d-e). The DC modulation could have positive implications for anti-tumor immunity, since the predominant role of DCs is to activate T cells, while macrophages clear apoptotic cells and microbes through phagocytosis. Therefore, both the metformin-induced transcriptomic activation of dendritic cell activation markers and increase in the proportion of DCs within the APCs were consistent with an idea that metformin could play a role in human dendritic cell activation, a notion which has not been made prior to the present study. The remaining APCs, consisting of monocytes and macrophages referred to as mono-macs (CD11c+, HLA-DR+, CD14+), also exhibit a significant decrease in markers associated with immunosuppressive M2 macrophages (Fig 4d,f) in corroboration to previous mouse studies and human in vitro co-culture studies^43–45^, which we now extend to authentic ex vivo human tumor tissue cultures.

To explore the effects of metformin in vivo, we investigated Wap-Myc tumor samples from metformin-treated mice for conventional dendritic cell (CD103+) and CD8 T cell (CD8+) numbers but found no significant changes in the cells expressing either marker (Fig 4g-i). These findings were consistent with our findings in PDEC-TIME, where there was an increase in APC activation, but not in overall numbers. Spatial 10x genomics profiling of Wap-Myc mouse tumor tissue from metformin treated animals revealed enriched dendritic cell activation in metformin-treated mouse compared to control mouse material as determined by the gene expression profiles; metformin increased *Irf7, Bst2, and Rsad2* expression, while macrophage marker, *CD68*, expression was downmodulated (Fig 4j-m; **supplementary Fig 4f-h**). Details of how the 10x genomics and how the tumor areas were classified are explained in supplementary **(Supplementary Fig 5a).** The results from human explants and mouse tumor tissue, treated with venetoclax and metformin (alone or in combination) together suggest that specifically metformin, and not venetoclax, affects dendritic cell activation, without altering their numbers.

### The mitochondrial respiratory complex I regulates DC activation and DC mediated activation of CD4+ T cell proliferation

To determine whether metformin activates APCs directly or through heterotypic cellular interactions, we isolated monocytes from human peripheral blood mononuclear cells (PBMC) of healthy donors. The PBMC-derived monocytes were then differentiated into either macrophages or dendritic cells. Flow cytometric analysis showed that metformin skewed macrophage polarization towards a less immunosuppressive subtype, as indicated by a decrease in CD163 and CD206 surface markers commonly associated with immunosuppressive macrophages, corroborating publications with similar findings (Fig 5a-d; ^44–46)^. RNA bulk sequencing of metformin-treated monocyte-derived DCs from 6 donors showed clear evidence for an increase in dendritic cell activation (Fig 5e, **supplementary 6a**). Additionally, flow cytometry analysis of monocyte-derived DCs revealed that treatments containing metformin induced a higher proportion of CD86-high, HLA-DR-high activated DCs from the total population (Fig 5f-g) suggesting that metformin indeed directly contributes to DC activation. Flow cytometry analysis of DC surface activation markers, and a transcriptional increase in dendritic cell activation gene sets suggests a less documented role of dendritic cells as a potential mediator of metformin’s anti-tumor effect in humans.

**Fig. 5.**
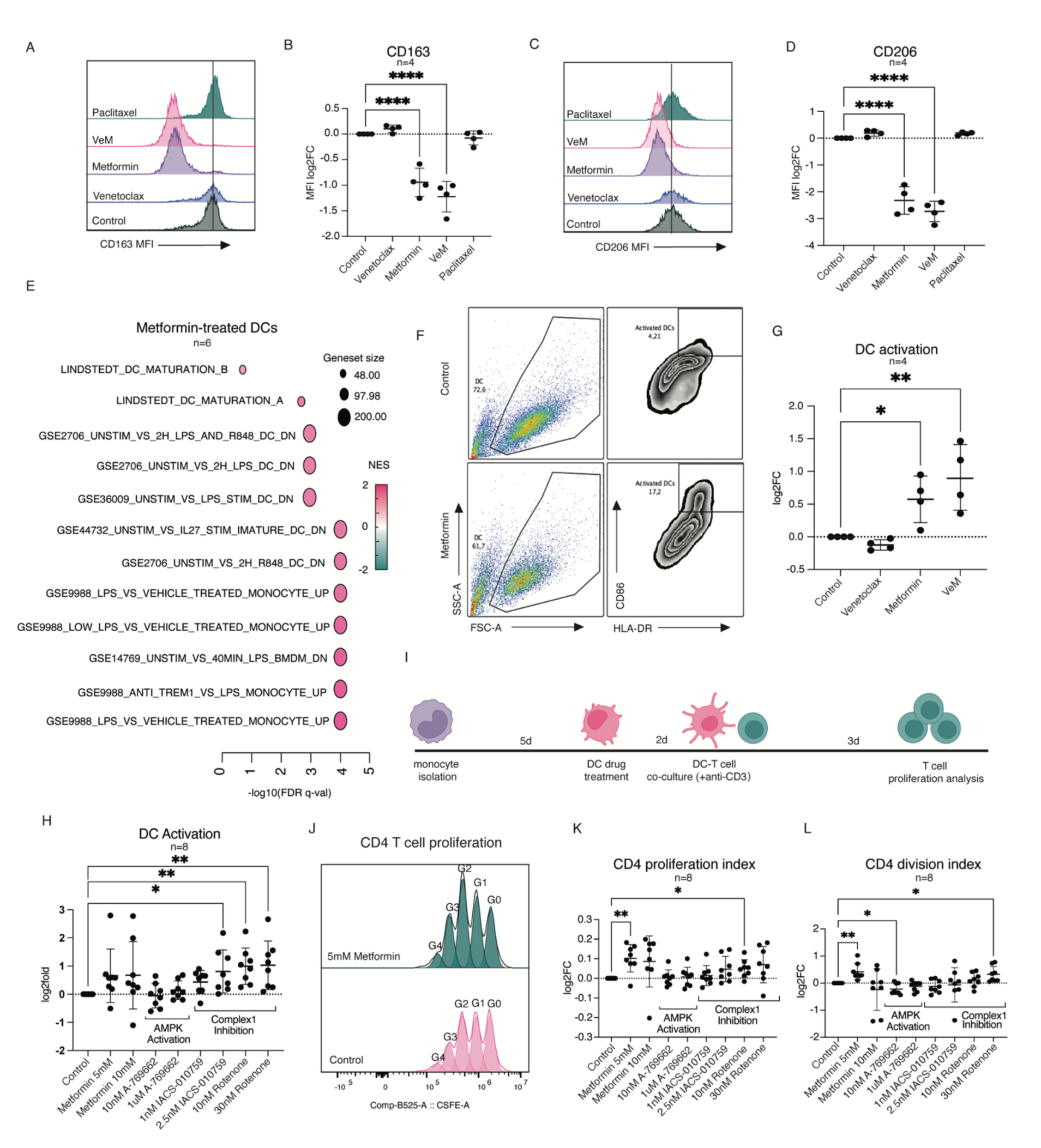
Complex 1 inhibition contributes to DC activation and CD4+ T cell proliferation. **a,** representative flow cytometry histogram of median fluorescence intensity of CD163 of monocyte-derived macrophages following treatment. **b,** Quantification of (a) from monocyte-derived macrophages of 4 biologically independent PBMC donors. **c,** representative flow cytometry histogram of median fluorescence intensity of CD206 of monocyte-derived macrophages following treatment. **d,** Quantification of (c) from monocyte-derived macrophages of 4 biologically independent PBMC donors. **e,** GSEA comparison of differentially expressed genes between 10mM metformin and control treated PBMC-derived dendritic cells from n=6 biologically independent PBMC donors. **f,** flow cytometry gating guide of HLA-DR and CD86 high-expressing DCs which are referred to as “activated DCs.” **g,** Quantification of the percentage of activated DCs following treatment from n=4 monocyte-derived DC samples with significant increase in surface expression of CD86 and HLA-DR following metformin (p=0.0226) and VeM (p=0.0014) treatments **h,** Quantification of DC activation following a larger panel of treatments with significant DC activation following 2.5nM IACS-010759 (p=0.0287), 10nM rotenone (p=0.0105), and 30nM rotenone (p=0.0309). **i,** schematic of timeline for DC, T cell co-culture assay. **j,** Representative T cell proliferation plot. G0 refers to the undivided generation, with each increasing G number referring to the number of proliferative cycles **k,** flow cytometry analysis of CD4+ T cell proliferation index indicating the proportion of the proliferating samples that keep proliferating (G1-G4) with significant increases in CD4+ T cell proliferation following 5mM metformin (p= 0.0043) and 10nM rotenone (p=0.0161) treatments., **l,** flow cytometry analysis of CD4+ T cell division index indicating the proportion of the total sample that is dividing (G0-G4) with significant increase in overall CD4+ T cell division following 5mM metformin (p= 0.0050), and 30nM rotenone (p=0.0148) treatments, and a significance decrease in CD4+ T cell proliferation following 10nM A769662 treatment (p=0.0184). All data are presented as mean values +/− SD. Statistical significance was tested with a one-way ANOVA with Fishers exact test.

Metformin is a medicinal biguanide, which acts as a weak inhibitor of mitochondrial respiratory complex I^47^. Inhibition of complex I decreases generation of ATP from oxidative phosphorylation and the resulting increase in AMP:ATP and ADP:ATP ratios activates adenosine 5′-monophosphate kinase (AMPK), an event commonly observed in metformin-treated cells in vitro and in vivo^48, 49^. To test whether metformin induces DCs activation through activation of AMPK or inhibition of the mitochondrial complex I, an experiment was designed to evaluate separately the effect of AMPK-activation, and complex I inhibition on DC activation. We found that inhibition of mitochondrial complex I with metformin, with the classical complex I inhibitor rotenone or with a specific ubiquinone reduction site targeted complex I inhibitor IACS-010759^50^ led to an increase in dendritic cell activation, whereas direct pharmacological allosteric AMPK activator A-769662 had no noticeable effect (Fig 5h). These results suggest that the mechanism by which metformin activates dendritic cells is largely through complex I inhibition.

Dendritic cells which express elevated levels of activating co-stimulatory signals, including CD86, promote T cell proliferation and acquisition of their cytotoxic abilities^51^. Thus, we explored whether metformin’s effect on DC activation has functional consequences on the activation of T cells. We pre-treated dendritic cells with metformin, A-769662, IACS-010759 and rotenone for 48hrs, and then co-cultured the DCs with autologous T cells in the absence of drugs. Importantly, we used anti-CD3 in the co-culture to fulfil the requirement of T cells to not only receive secondary co-stimulatory signals from APCs, but also to have the primary signal through the T cell receptor which we were supplementing artificially (Fig 5i). An increase in CD4+ T cell proliferation was quantified as a shift in the proportion of T cells in proliferation (G0 = unproliferated T cells, G1 = first division, etc.), or in other words, the number of divisions of already-proliferating cells (Fig 5j). This “proliferation index” clearly showed that the trend in CD4+ T helper cell proliferation mirrored the trend in dendritic cell activation (Fig 5k). Meanwhile, the “division index”, which is a measure of the total proportion of cells that begin to divide (the ratio of G1-4 to G0), revealed a weak yet consistent trend, with a significant negative impact on overall CD4+ T cell proliferation following co-culture with DCs treated with AMPK-activator, A-769662 (Fig 5l). This can be a result of AMPK-activation interfering with DC maturation as seen in mice^52^. These data suggest that complex I inhibition may improve an antigen-specific immune response through the activation of DCs.

## DISCUSSION

The importance of targeting tumor cells and exploiting the TIME simultaneously is evident by the increasing number of clinical trials combining targeted therapies or chemotherapies with immunomodulatory compounds^53^. However, microenvironmental interactions and heterogeneity within the tumor microenvironment that affect treatment response^54–56^, can be difficult to model in the laboratory. In vivo animal models may require surrogate, species-specific reagents with different pharmacological properties than the original therapeutic drug. In vitro organoid models capture disease heterogeneity and tumor intrinsic features to some extent but are lacking most immune cell components^57–59^. Some organoid models are being adapted, for example, to utilize autologous primary tumor-specific CD8+ T cells to screen tumor cell killing in response to increased drug-induced T cell cytotoxicity^60^. Although beneficial in terms of throughput, this is limited to CD8+ T cell-mediated cytotoxicity. Moreover, utilizing the tumor itself is crucial because anti-tumor T cell responses can be tumor site specific, even within the same patient^61, 62^. So, while peripheral blood can be powerful as a predictor of monitoring clinical outcome^63^, tumor resident immune cells can be more informative for drug development by capturing the complex tumor-TIME dynamics which can shaped by the baseline immune composition^64^. Indeed, there is a clear correlation with the full TIME repertoire and clinical outcome, prompting more attention to models with a more diverse repertoire of autologous tumor immune cells^65–68^.

Here we explored the opportunities provided by breast cancer patient-derived explant cultures (PDECs). While organoids can be considered as “reconstruction” models where tumors are dissociated into single cell components that form new structures, explants are a “deconstruction” of the tumor which captures the original architecture, heterogeneity, and immune cell composition of the tumor – but in smaller pieces. Earlier works show that human ex vivo explants from various cancer types retain tumor and stroma components in addition to autologous immune cell populations and can respond to anti-PD-1 depending on the cancer type^35, 69, 70^. We deduced through single cell sequencing, gene expression profiling, flow cytometry, and cytokine profiling of PDECs in comparison to primary tumor material that PDECs also retain all major immune cell subtypes and baseline immune cell activity up to one week in culture, which makes these PDEC-TIME cultures an attractive model to explore tumor-TIME dynamics.

While we failed to see an increase in T cell activity or tumor cell death in breast cancer PDECs in response to either anti-PD-1 or anti-PD-L1, we did see strong immune activation in response to artificial T cell activation using an anti-CD3/CD28/CD2 tetramer similarly to Voabil (2021), suggesting that T cells within breast tumors lack tumor antigen reactivity. In response to anti-CD3/CD28/CD2, we first observed cell death appearing in PDECs with CellTox staining and an increase in immune activation markers IFNɣ, PRF1, and GZMB via qPCR from bulk RNA. Subsequent flow cytometry and cytokine profiling revealed that the cell death was specifically affecting CD45-tumor cells, while cytokines of T cell activation (e.g. IL-2, IFNɣ, IP-10) were increased along with T cell surface expression of checkpoint molecules like PD-1, PD-L1, PD-L2, and LAG-3. Altogether, these findings suggest that with the correct treatment strategy, breast tumor-resident immune cells in PDECs have the potential to be exploited for preclinical anti-tumor activity.

In our previous work on MYC-driven mouse models of breast cancer, we observed exceptionally durable tumor growth control following a triple-treatment of venetoclax+metformin (VeM), and anti-PD-1^27^. The ex vivo PDEC model, however, revealed a downregulation of all 27 tested cytokines following VeM treatment. Additional flow cytometry data revealed severe T-cell toxicity, especially among CD8+ T cytotoxic cells. We attributed this phenomenon to corroborating lymphotoxic effects of venetoclax on human T cells observed in vitro and in vivo^34, 71^. Nevertheless, gene expression profiling of a larger repertoire of immune cells revealed that other immune cell subsets were unaffected by VeM in terms of numbers, and that there was a significant increase in genes and cytokines suggestive of APC-activation. The survival and activation of APCs, including macrophages and dendritic cells, is highly interesting since APCs are known to mold the pro/anti-tumor properties of TIME, and thus plays a role in the efficacy of immune-checkpoint blockade^72, 73^.

Classically activated “M1-like” proinflammatory macrophages have been suggested as a standalone therapeutic strategy for breast cancer due to their tumoricidal properties^74^, and a biomarker of increased survival in response to trastuzumab^75^. Meanwhile, alternatively activated “M2-like” macrophages have been linked to poor prognosis in breast cancer. Metformin has been shown to prevent M2 polarization of macrophages, induce M1 polarization of macrophages, and increase the efficacy of checkpoint inhibition in human and mouse studies^44–46, 76, 77^. Extending the previous results, we demonstrate in PDECs that 5mM-10mM of metformin directly decreases the proportion of immunosuppressive CD163+ M2-like macrophages.

Dendritic cells are highly potent APCs which can trigger robust, antigen-specific T cell activation in vivo upon maturation, and their increased presence is generally considered a good prognostic marker in breast cancer^78, 79^. Our data show that while metformin does not influence the total number of APCs, it induces a higher proportion of activated DCs in PDECs, and we also observed that metformin promotes DC activation of isolated DCs from human PBMCs. Thus, our data suggests that metformin has a direct DC-activating function, contrary to the idea that DC activation was solely induced through the release of tumor antigens following VeM treatment.

Metformin acts as a weak inhibitor of mitochondrial respiratory complex I^47^ and as an AMPK-activator^48, 49^. To investigate how metformin mediates its DC activating function, we explored these two most well-known mechanisms of metformin action. We only observed that the metformin effects on DC activation were phenocopied by respiratory complex I inhibition (CI-i), but not by direct allosteric activation of AMPK. Previous reports have suggested that CI-i of dendritic cell precursor monocytes with rotenone inhibits the development of monocytes into immature dendritic cells^80^, however, to our knowledge metformin’s DC-activating effects mediated via CI-i, shown here, have not been previously reported. If metformin inhibits the differentiation of monocytes to immature DC’s, but also simultaneously promotes DC activation, then what would be the net effect of metformin on the fully physiological TIME of breast cancer? Since evidence indicates that resident DCs within breast tissue are already in an immature DC state^81^, we propose that CI-i would immediately and beneficially target the tumor resident DCs by increasing tumor-antigen presentation by now-mature DCs. Therefore, adding metformin as an agent to explorative or standard cytotoxic treatments could leverage the tumor-antigen-releasing effects of the cytotoxic component, through simultaneously promoting DC-activation-a concept that should be especially considered in the scope of combination immunotherapies aimed at maintaining drug-induced T cell responses.

Mechanistically, pharmacological inhibition of CI has a variety of different cell energy metabolism and oxidative stress related effects, which alone or via additive or synergistic effects could explain the DC activation. While the exact mechanism how CI-i induces DC maturation remains outside of the scope of present study, we note that previous study has revealed an increase in DC maturation and subsequent T cell activation in response to free oxygen radicals^82^, which are in some circumstances released in response to respiratory CI inhibition. As an alternative to the causal role of free oxygen radicals, the DCs may undergo a “metabolic shift” in response to CI-i treatment^83–86^. Dendritic cells undergo a metabolic shift from oxidative phosphorylation to aerobic glycolysis in response to toll-like receptor agonists leading to DC maturation^52^. As metformin and CI-i decrease oxidative phosphorylation^87^, it is not inconceivable for a compensatory pathway to take over leading to DC maturation.

In summary, we demonstrate PDEC as a versatile preclinical immuno-oncology model of human tumor immune microenvironment that can be easily adapted for a variety of research techniques like single-cell sequencing, and other studies which require the extraction of viable, single cells. Using the PDEC-TIME model, we reveal a hitherto unknown role of metformin for maturation of antigen presenting cells, specifically dendritic cells, highlighting the potential clinical translatability of complex 1 inhibition as means of boosting immunotherapy treatments.

## MATERIALS AND METHODS

### Cell Lines and Reagents

Human peripheral blood mononuclear cells (PBMCs) were cultured in RPMI medium supplemented with 10% heat-inactivated FBS (Biowest), 100 U penicillin-streptomycin (Gibco), and 2 mM L-Glutamine (Gibco). PDECs were cultured in MammoCult (StemCell technologies), and the MammoCult media was supplemented with MammoCult proliferation supplement #05622 (StemCell technologies), 20 μg/mL gentamicin (Sigma), 0.1 μg/mL amphotericin B (Biowest) and 10,000 U/mL penicillin/streptomycin (Lonza). Cells and PDECs were grown in a humidified incubator at 37◦C under 5% CO2, and atmospheric oxygen levels.

PDECs were treated with 25ul/mL anti-CD3/CD28/CD2 (Stemcell Technologies), 100ug/mL Atezolizumab (Selleck Chemicals), 50ug/mL Pembrolizumab (MedChem), 10-100nM Venetoclax (MedChem Express), 5-10mM Metformin (MedChem Express), 10-50nM Paclitaxel (MedChem Express), 1-2.5nM IACS-010759 (Selleck Chemicals), 10-30nM Rotenone (Sigma-Aldrich), and 10nM-1uM A-769662 (Sigma-Aldrich), 100ng/ml LPS.

### Isolation of Biological Material and Three-Dimensional (3D) Culture

Fresh tissue was obtained from the elective breast cancer surgeries performed at the Helsinki University Central Hospital **(Supplementary Fig 7a-b**) (Ethical permit: 243/13/03/02/2013/ TMK02 157 and HUS/2697/2019 approved by the Helsinki University Hospital Ethical Committee). Patients participated in the study by signing an informed consent form. Tissues were collected from tumors. From each tumor, a portion was taken for immunohistochemical, a second portion was frozen at −80°C DNA/RNA/protein analysis, and the reminder was used for the 3D cultures. Explants were produced by incubating the samples overnight in collagenase A (3 mg/ml; Sigma) containing MammoCult media (StemCell technologies) with gentle shaking (130 rpm) at +37°C. The resulting explants were collected via centrifugation at 353 rcf for 5 min and washed once with 1xPBS. Isolated explants were embedded in Cultrex Reduced Growth Factor Basement Membrane Extract, Type 2 (R&D Systems) and plated on 8-chamber slides (Thermo Scientific).

### Flow Cytometry

Explants were harvested by washing the wells twice with 1x PBS, then resuspended in 400ul +4°C Cultrex Organoid Harvesting Solution (Bio-Techne Sales Corp.) and incubated at +4°C with gentle shaking for 30min. Samples were pipetted onto Falcon round-bottom tubes with cell strainer caps (Corning) and centrifuged at 400rcf for 5 minutes at +4°C. Samples were resuspended in 100ul of flow cytometry staining buffer (1xPBS, 10% heat-inactivated FBS (Gibco)) and the appropriate antibodies (Supplementary data 2) for 45minutes at +4°C in the dark. Samples were washed twice with flow cytometry running buffer (1xPBS, 1% heat-inactivated FBS (Gibco), and resuspended in running buffer for analysis.

Samples sorted with BD Influx or Sony SH800Z, or analyzed using BD FACSAria II, or NovoCyte Quanteon (Biomedicum Flow Cytometry Unit). Analysis was done using FlowJo v10.8.1, and graphs were generated using GraphPad Prism v9.

Flow cytometry antibodies listed in **Supplementary table 1**.

### Macrophage Polarization Assay

CD14+ monocytes were isolated using positive magnetic separation with manufacturer instructions (Miltneyi) and were cultured in Iscove’s modified Dulbecco’s medium (IMDM; Thermo Fisher) supplemented with 10% heat-inactivated FBS (Gibco), penicillin/streptomycin, and 50ng/mL human M-CSF (Miltenyi Biotec), and 50ng/mL Human IL-4 (Miltenyi Biotec). Monocytes were plated into 6-well plates (5×10^5^ cells) and incubated for 6 days with one medium change. Media from differentiated macrophages was replaced with fresh media containing drugs. The adherent macrophages were detached using macrophage detachment solution DXF (Sigma-Aldrich) for 40 min at +4°C, washed with flow cytometry running buffer, and blocked with Fc-blocking antibody (eBioscience) in flow cytometry running buffer for ten minutes at room temperature (according to manufacturer instructions) before adding fluorescent antibodies. Macrophage Median Fluorescence Intensity of surface CD163 and CD206 expression was analyzed on the Novocyte Quanteon flow cytometer.

### Dendritic cell activation assay

CD14+ monocytes were isolated using positive magnetic separation according to manufacturer protocol (Miltneyi Biotec) and cultured in RPMI medium supplemented with 10% heat-inactivated FBS (Biowest), 100 U penicillin-streptomycin (Gibco), and 2 mM L-Glutamine (Gibco) with added 100ng/mL Human GM-CSF (Miltenyi Biotec), and 100ng/mL Human IL-4 (Miltenyi Biotec) for 5 days only adding media once in between. On day 5, DCs were harvested and plated on a 96-well flat-bottom plate. Samples were incubated with 200ul fresh media containing drugs for 48hrs. Dendritic cells were harvested and analyzed using flow cytometry on novocyte Quanteon. Dendritic cells expressing high levels of HLA-DR and CD86 were considered activated.

### T cell proliferation assay

T cells were isolated using Pan T magnetic isolation beads (Miltenyi Biotec) according to manufacturer protocol, and stained with 1uM CSFE in 1xPBS in 37°C 5% CO_2_ for 10 minutes before washing the samples with media. Stained T cells were co-cultured with autologous dendritic cells at a ratio of 5:1 (T: DC) in dendritic cell media. 100ng/ml of Ultra-Leaf purified anti-human CD3 (Biolegend 300413) was added to each sample, and anti-CD3/CD28/CD2, Immunocult (StemCell Tech) was used as a positive control. Samples were incubated for 72hrs at 37°C 5% CO_2_ before being harvested and analyzed using flow cytometry. T cell proliferation was quantified as the division index and proliferation index as calculated by FlowJo v10.8.1 software.

Division Index: Total Number of Divisions / The number of cells at start of culture Proliferation Index: Total Number of Divisions / Cells that went into division

### scRNA Sequencing and analysis

PDEC samples were harvested and stained with CD45+ to isolate CD45+ leukocytes with fluorescence-activated cell sorting from the total tumor tissue. Up to 30,000 CD45+ cells were collected for single cell 3’ v3 library preparation with 10X genomics Chromium. These single cell libraries were sequenced on an Illumina NovaSeq6000. Raw data were demultiplexed, aligned to GRCh38-2020-A and gene count data were generated by CellRanger (cellranger-4).

Raw count matrices were further analyzed in Seurat. Barcodes with <20% mtRNA,> 400 unique read-counts and number of features between 200 and 6000 were retained as cells.

Next the data were integrated using Seurat’s Canonical-Correlation-Analysis (CCA)^88^, clustered and visualized by a UMAP. Celltypes were identified based on established marker genes. Significantly enriched cell types between cultured and tissue samples were calculated using the MASC algorithm^89^. The copy-number variation was assessed with inferCNV (https://github.com/broadinstitute/inferCNV) to identify the clusters of cancer cells.

### 3’ RNA Sequencing

Total RNA was isolated using RNeasy Plus (Qiagen) which contains a gDNA eliminator column. RNA sequencing libraries were prepared from 100 ng of total RNA using either the ScriptSeq Complete Gold Kit or the NEBNext Ultra Directional RNA Library Prep Kit for Illumina depending on the RNA integrity. Using the ScriptSeq Complete Gold Kit, the ribosomal RNA was removed first from the total RNA using the Ribo-Zero™ Gold rRNA Removal Kit after which the RNA was fragmented chemically. The libraries were prepared according to the manufacturer’s instructions. Finally, the library was assessed with the Agilent Bioanalyzer.

The NEBNext Ultra Directional RNA Library Prep Kit for Illumina was used to generate the cDNA libraries for next generation sequencing. First, the ribosomal RNA depleted samples (10 ng) were fragmented to generate the inserts around 200 bp. The libraries were prepared according to the manufacturer’s instructions. The library quality was assessed with Bioanalyzer (Agilent DNA High Sensitivity chip) and the library quantity with the Qubit (Invitrogen).

Samples were sequenced with the NextSeq 500—Illumina instrument using 75 PE reads with a sequencing depth of 33 M reads/sample. Differentially expressed genes between different groups were found using state-of-the-art statistical methods and packages, such as edgeR/DESeq2. The Gene Set Enrichment Analysis 3.0 (Broad Institute) was used to analyze the differences in the gene expression profiles. GSEA results were visualized using GraphPad Prism v.9.5.0.

### Mouse Tissue Spatial Transcriptomics

Two cryopreserved tumor tissue sections from one metformin treated and one untreated Wap-Myc mice were profiled for spatial transcriptomics using the 10x Genomics Visium spatial RNA-sequencing technology with a resolution of 55 µm per spot. The tissues were cryosectioned at 10-µm thickness onto the Visium library preparation slide, fixed in methanol for 30 min, stained with Hematoxylin and Eosin Stain Kit (Vector Laboratories) and stored at −20℃ until library preparation. The slide was imaged using Zeiss Axio Imager.

Sequencing library preparation was performed according to the Visium Spatial Gene Expression user guide (CG000239 RevC, 10x Genomics) using a 12-min tissue-permeabilization time that was tested earlier with the Tissue Optimization Slide to be optimal for the mouse tumor tissues. Libraries were sequenced with Illumina NovaSeq6000 sequencer at FIMM Genomics core facility at University of Helsinki, with the aimed depth of 75 million reads per section (corresponding approximately 50.000 reads per tissue covered array spot).

The analysis was done using R version 4.2.2 and Seurat version 4.3.0. In order to maintain only good quality spots and relevant spots in analysis, spots with lesser than 500 Features or more than 25% of mitochondrial transcripts or more than 1% of hb transcripts were excluded from the analysis. Data was normalized using SCTransform^90^ and afterwards, principal component analysis and dimension reduction with UMAP^91^ was done with default parameters and with Seurat functions accordingly. Identification of clusters was done with Seurat FindClusters function and resolution of 0.8. Top significantly (padj<0.05) differentially expressed genes **(supplementary Fig 5a)** obtained from Seurat FindMarkers function (Wilcox rank sum test) were used for cluster annotation. Differentially expressed genes between different tumor areas were defined by using the same function and Wilcox ran sum test.

### Nanostring gene expression profiling

Total RNA was isolated using RNeasy (Qiagen), and the DNAase removal step was performed after the isolation (Zymo research). Samples went through QC (Quibit), Gene expression analysis was conducted on the NanoString nCounter gene expression platform (NanoString Technologies). Due to the systemic reduction of TILs in explants, the samples were normalized to the genes that create the TIL score: T cell, CD45, B cell, Cytotoxic cell, and macrophage genes excluding those with counts below 100 (*CD3G, CD3D, CD3E, SH2D1A, CD6, PTPRC, BLK, MS4A1, TNGFRSF17, CD19, CD84, CD68, CD163, CTSW, KLRB1, KLRD1, GzMB, PRF1, GZMA, GNLY, KLRK1. GZMH, CD8, CD8A, CD8B, CD4*) before comparing activity profiles. Significance was calculated with nSolver’s directional significance score calculated as t-statistic for each gene against each covariate in the model, taking the sign of the t-statistics into account.

nCounter Human PanCancer Immune Profiling Panel consisting of 770 genes from different immune cell types, common checkpoint inhibitors, CT antigens, and genes covering both the adaptive and innate immune response. Per sample, 50 ng of total RNA in a final volume of 5 μl was mixed with a reporter codeset, and hybridization buffer, and capture codeset. Samples were hybridized overnight at 65°C for 20 hours. Hybridized samples were run on the NanoString nCounter SPRINT profiler.

### Multiplex and Standard Immunohistochemistry

Tissues and explant cultures were fixed with 4% paraformaldehyde (PFA) and embedded in paraffin. The samples were sectioned into 5 μm slices and deparaffinized. The heat-induced antigen retrieval was performed with a microwave oven or a pressure cooker in a citrate buffer solution (Dako). Histochemical stainings were carried out using standard techniques for IHC and IHC-IF. Images were taken with a Leica DM LB microscope or with a Zeiss AxioImager 1 (Biomedicum Imaging Unit, University of Helsinki). Multiplex images were stained in two rounds and tissue sections were scanned on the Zeiss AxioImager Z1 scanner at FL20X.

The list of used antibodies is shown in **Supplementary Table 2**.

### Immunofluorescent staining

Three-dimensional (3D) cultured breast cancer explants were fixed with 4% paraformaldehyde for 15 min at room temperature and washed three times with PBS. The tissue explants were permeabilized with 0.5% Triton X-100 in PBS for 10 min at RT and blocked in an IF buffer (0.1% BSA, 0.2% Triton X-100, 7.7 mM NaN3, and 0.05% Tween 20 in PBS) supplemented with 10% (v/v) normal goat serum for 1 h. Explants were then incubated with the primary antibody diluted in a blocking solution overnight at 4 °C. Following incubation, explants were washed three times with an IF buffer and then incubated using the appropriate Alexa Fluor secondary antibody diluted in an IF buffer with 10% goat serum. After 60 min of incubation at RT, the explants were washed with an IF buffer as before and the nuclei were counterstained with Hoechst 33258 (Sigma). Instead of antibodies, cell death staining was done with the CellTox green (Promega) at a dilution of 0.25ul CellTox/500ul media for 20 min. Samples were then washed twice with 1xPBS, fixed with 2%PFA for 20min, washed twice with 1xPBS, and stored in +4°C until imaging. Slides containing tissue explants were mounted with the ImmuMount reagent (Fisher Scientific). Images of the structures were acquired using a Leica TCS SP8 CARS confocal microscope using an HC PL APO CS2 40x objective (Biomedicum Imaging Unit, University of Helsinki).

### Cytokine profiling

PDEC cytokine secretion was analyzed from cleared PDEC culture supernatants using Bio-Plex Pro Human Cytokine 27-plex assay kit (Bio-Rad, cat. M500KCAF0Y) and Bio-Plex 200 System (Bio-Rad) according to the manufacturer<u>’</u>s instructions. Results were analyzed using Bio-Plex Manager 6.0 software (Bio-Rad Laboratories).

Cytokines with >10% of datapoints outside the detection range were excluded from the analyses. Remaining values lower than the detection limit were replaced by 0.5× lowest measured value. Further data analyses and visualizations were performed using R (v.4.0.4^92^, tidyverse v.1.3.1). To identify changes in untreated PDEC cytokine secretion over time, we calculated log_2_FoldChanges between Day 6 and Day 3 cytokine levels and analyzed their deviance from zero (unchanged) with one-sample T-test. Resulting Benjamini-Hochberg-adjusted p-values and average log_2_FoldChanges were visualized as a volcano plot. For comparing different treatments, we used DMSO-treated PDEC cytokine levels as baseline and calculated log_2_FoldChanges for the indicated treatments. Unsupervised hierarchical clustering was performed with R (function hclust) based on log2FoldChanges and visualized as a heatmap using ComplexHeatmap package (v2.6.2)^93^. Bar graphs (median ± IQR) and line plots (median) were plotted using log_2_FoldChange values, and statistical significances reported as Benjamini-Hochberg adjusted p-values from one-sample Wilcoxon signed rank tests (deviance from zero).

### q-PCR analysis

Total RNA was isolated from cell lines and primary cell cultures using the Qiagen RNEasy Kit according to the manufacturer’s instructions, while the cDNA synthesis was performed with the Maxima First Strand cDNA Synthesis Kit for RT-qPCR (Thermo Scientific). Real-time RT-PCR was performed with LightCycler® 480 II (Roche) using DyNAmo ColorFlash SYBR Green (Thermo Scientific). The gene-specific primer sets were used at a final concentration of 0.2 mM Primers are listed in **Supplementary Table 3**.

### Statistical analysis

We report our results as the mean ± standard deviation (SD). Datasets were analyzed using Fischers exact test. All the experiments with representative images (Immunohistology, and immunofluorescence stainings) have been repeated at least thrice. When comparing multiple groups, the p values were calculated using one-way ANOVA, unless otherwise specified in the figure legend.

## Data availability

Source data for BRB sequencing, scSequencing, and gene expression profiling can be found deposited at 10.5281/zenodo.7906989, Relevant manuscript data are available from the authors upon request.

## ACKNOWLEDGEMENTS

We are grateful to the patients who participated in this research by donating breast tumor tissue samples and made it possible, and to the surgical personnel at Helsinki University Hospital who assisted with the recruitment and collection of the sample material. We are grateful to Biomedicum Imaging Unit (BIU), Biomedicum Functional Genomics Unit (FuGU), and Biomedicum Flow Cytometry Unit (from HiLIFE, University of Helsinki and Biocenter Finland) for their services. WI thank the Klefström laboratory personnel for discussions and critical comments on the manuscript. Schematic images were created with BioRender.com. We thank the Finnish Cancer Institute (FCI) for financial support. This work was supported by grants from the Academy of Finland, Business Finland, the Finnish Cancer Organizations, the Sigrid Juselius Foundation, Jane and Aatos Erkko Foundation, and RESCUER project, which has received funding from the European Union’s Horizon 2020 research and innovation programme under grant agreement No. 847912. This work was also supported by the Office of the Assistant Secretary of Defense for Health Affairs through the Breast Cancer Research Program under Award No. W81XWH2110773. Opinions, interpretations, conclusions, and recommendations are those of the author and are not necessarily endorsed by the Department of Defense. In addition, funds were received from Archimedes Foundation, Ida Montinin Foundation, Cancer Society of Finland, and iCAN Digital Precision Cancer Medicine Flagship.

## Notes

### Competing Interest Statement

The authors have declared no competing interest.

