## Supplementary Figures and Tables for "Respiratory Complex I Regulates Dendritic Cell Maturation in Explant Model of Human Tumor Immune Microenvironment"

Supplementary Fig 1

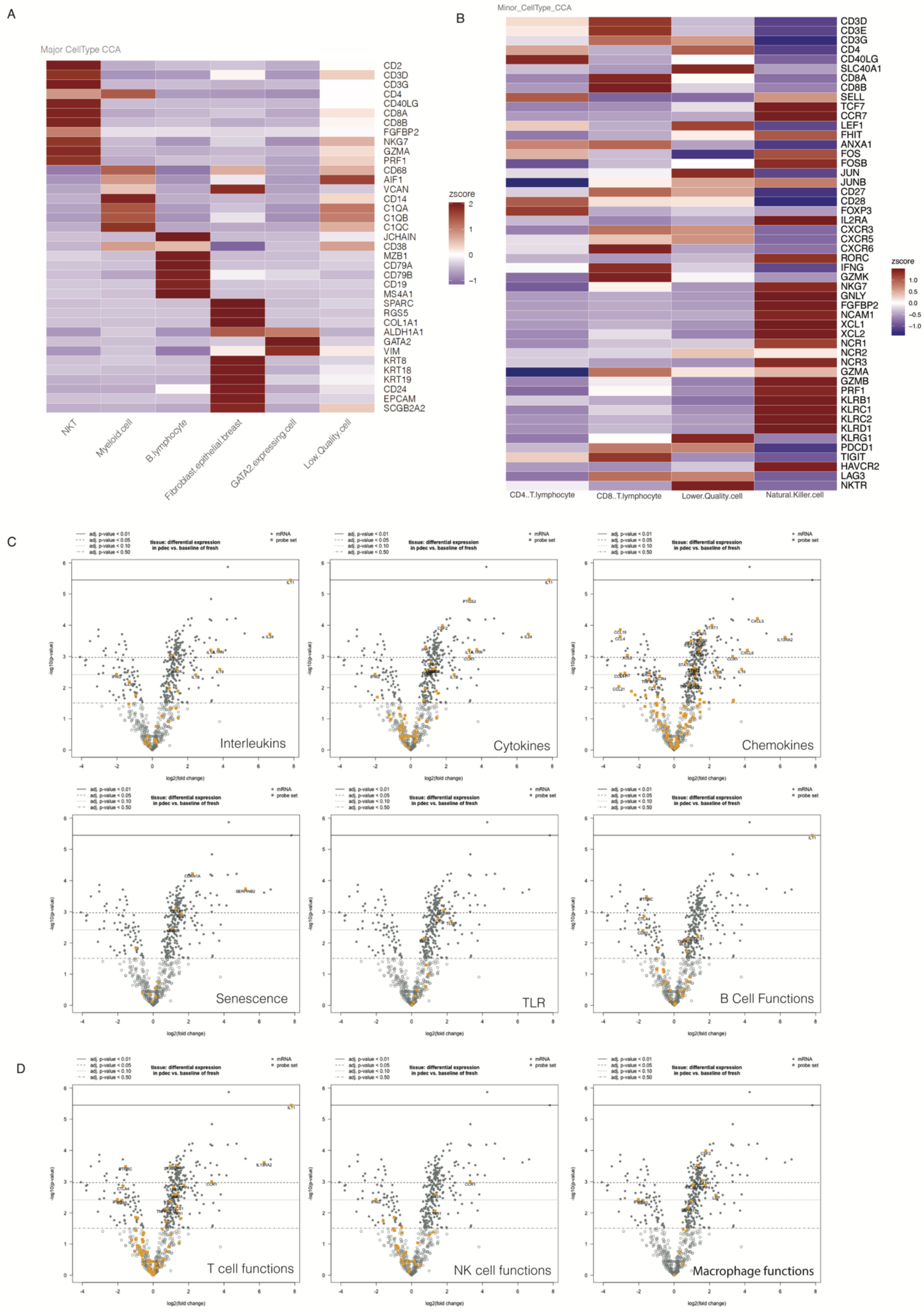

**Supplementary Fig. 1. Characterization of PDEC immune clusters and activity pathways a,** clustering of single cell RNA data into major and **b,** minor immune cell subtypes based on known gene expression profiles for each cell type **c,** volcano plots of the genes used to compare basal activity of pathways including NK cell functions, T cell functions, B cell functions, cytokines and interleukins, TLR, senescence and macrophage functions of PDECs grown 72hrs compared to primary tumor material. The significance of individual genes is separated by horizontal lines of defined adjusted p-values.

Supplementary Fig 2

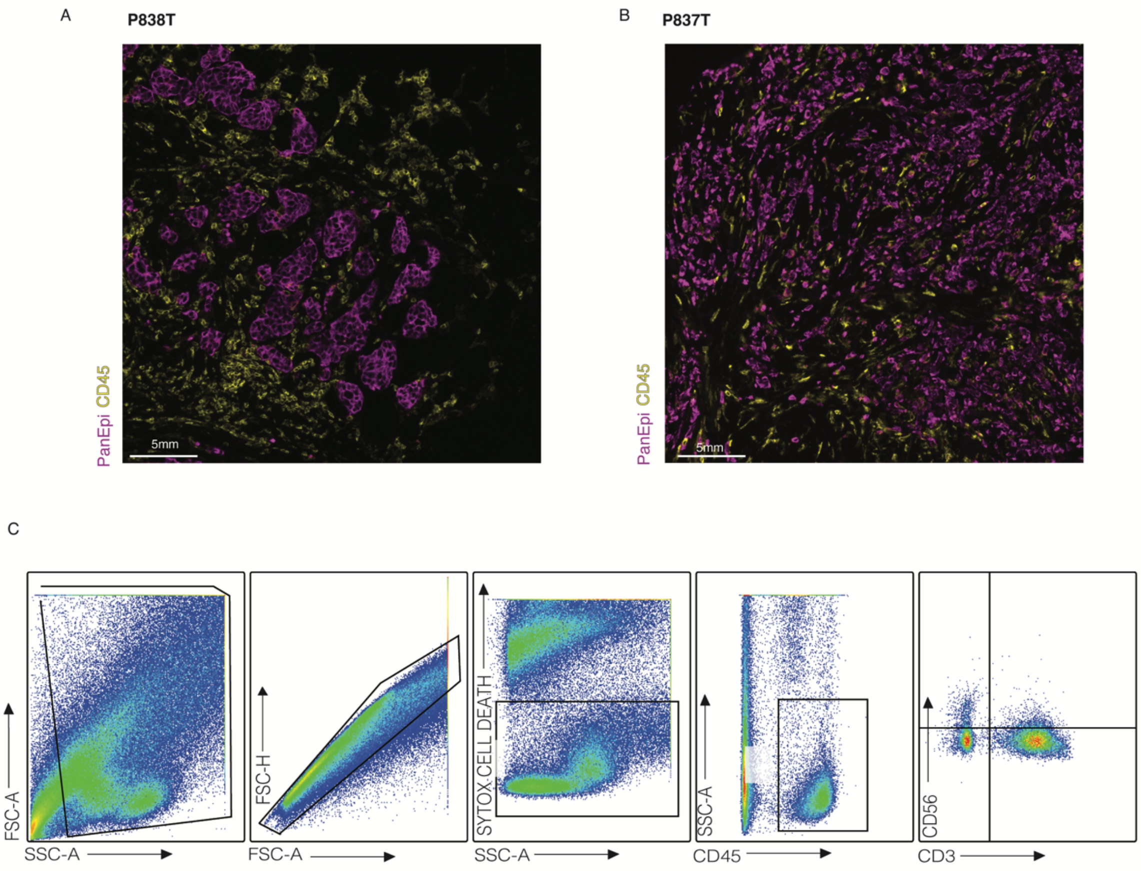


**Supplementary Fig. 2. Immune cell status of primary tumor material, and flow cytometry analysis of PDECs a,b,** Primary tumor immune infiltration of two randomly selected breast cancer samples corresponding to PDECs from figure. CD45+ leukocytes (yellow), tumor cells (purple). The scale is 5mm. P838T and P837T refer tumor samples of two biologically independent breast cancer sample donors. 2a. **c,** flow cytometry gating guide to determine checkpoint marker expression of CD3+ T cells from PDECs. Debris, doublets, and non-viable cells were removed from the analysis. CD45+ leukocytes and CD3+ CD56- T cells were analyzed for figure 2k-o.

Supplementary Fig 3

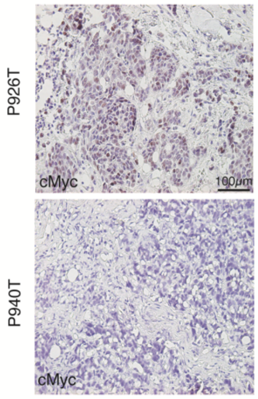


| | cMyc Staining |
|---|---|
| P899T | 0-20% |
| P900T | 61-100% |
| P905T | 0-20% |
| P910T | 0-20% |
| P916T | 0-20% |
| P917T | 41-60% |
| P924T | 61-100% |
| P926T | 61-100% |
| P939T | 21-40% |
| P940T | 0-20% |

**Supplementary Fig. 3. MYC protein staining of primary breast cancer tissue a,** c-MYC staining of 10 biologically independent patient tumors used in Fig 3b. Samples were scored for MYC positivity by from 0-100% in increments of 0-20%, 21-40%, 41-60%, and 61-100%.

Supplementary Fig 4

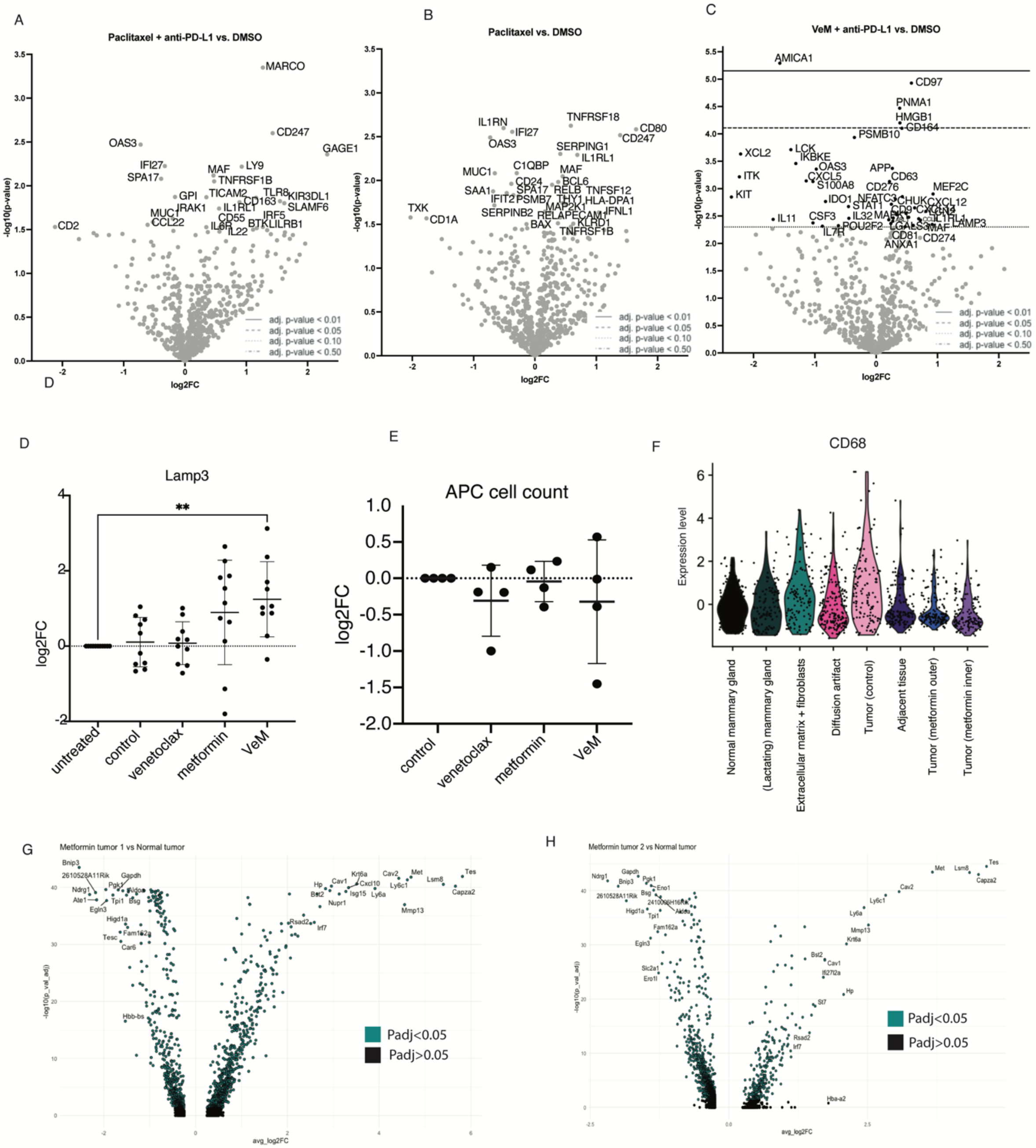


**Supplementary Fig. 4. Modulation of APCs in response to paclitaxel, VeM, and metformin-alone a,** volcano plot of gene-expression changes in response to paclitaxel + anti-PD-L1 **b,** paclitaxel and **c,** Venetoclax +metformin + anti- PD-L1 through nanostring gene expression profiling **d,** RT-qPCR increase of LAMP-3 in 10 PDECs in response to metformin **e,** flow cytometry of absolute number of APCs in 4 PDECs with no significant change in cell numbers **f,**

spatial transcriptomics of WapMYC mouse tumor tissue showing decreased CD68 macrophage expression (n=1) following metformin treatment **g,h,** volcano plot of differentially expressed genes in metformin treated tumor tissue

Supplementary Fig 5

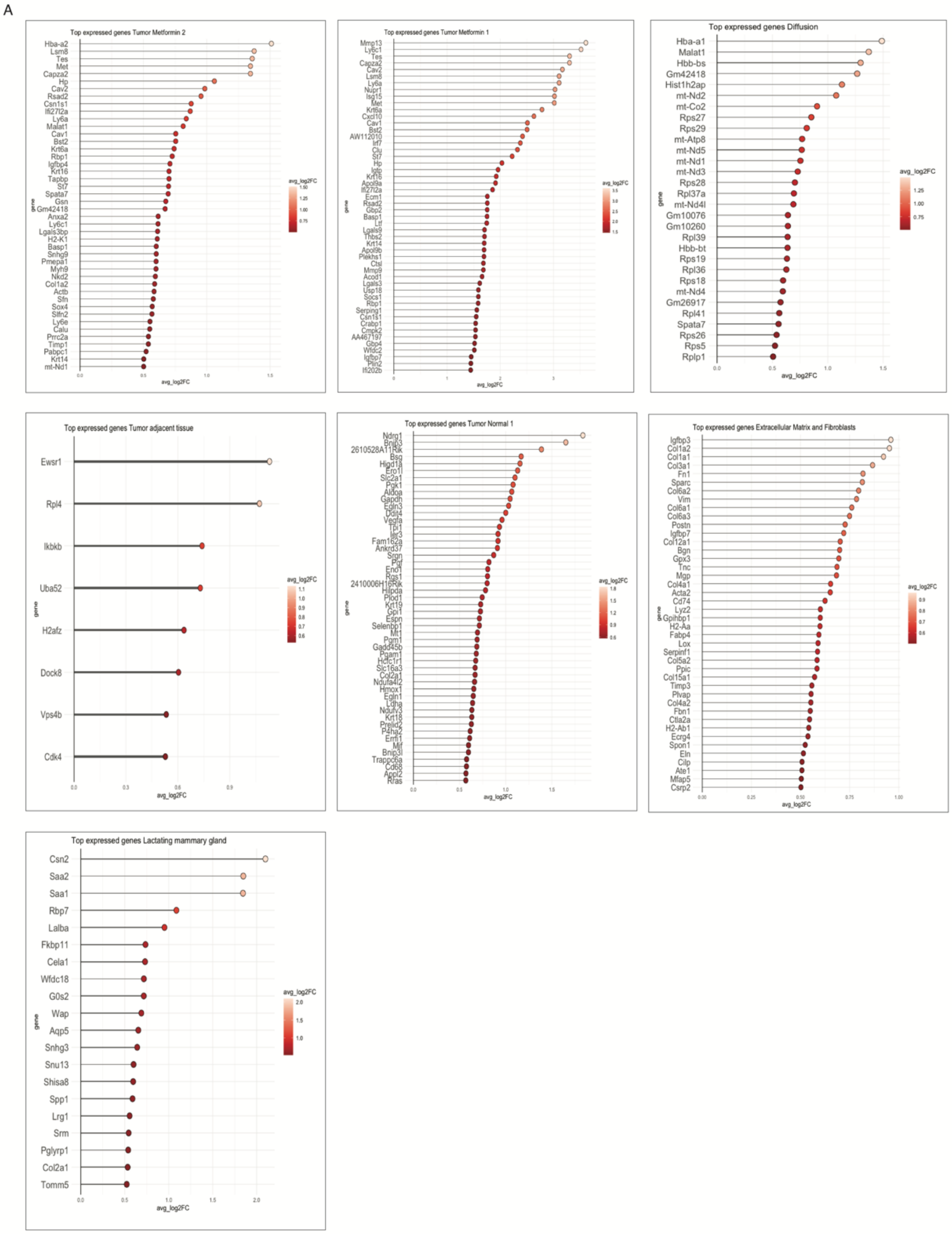


**Supplementary Fig 5. Characterization of tissue areas in spatial transcriptomics tumor tissue**

**a,** Top expressed genes used to characterize 9 independent tissue type clusters from spatial transcriptomics data for comparison from WAPMYC mouse tumor tissue

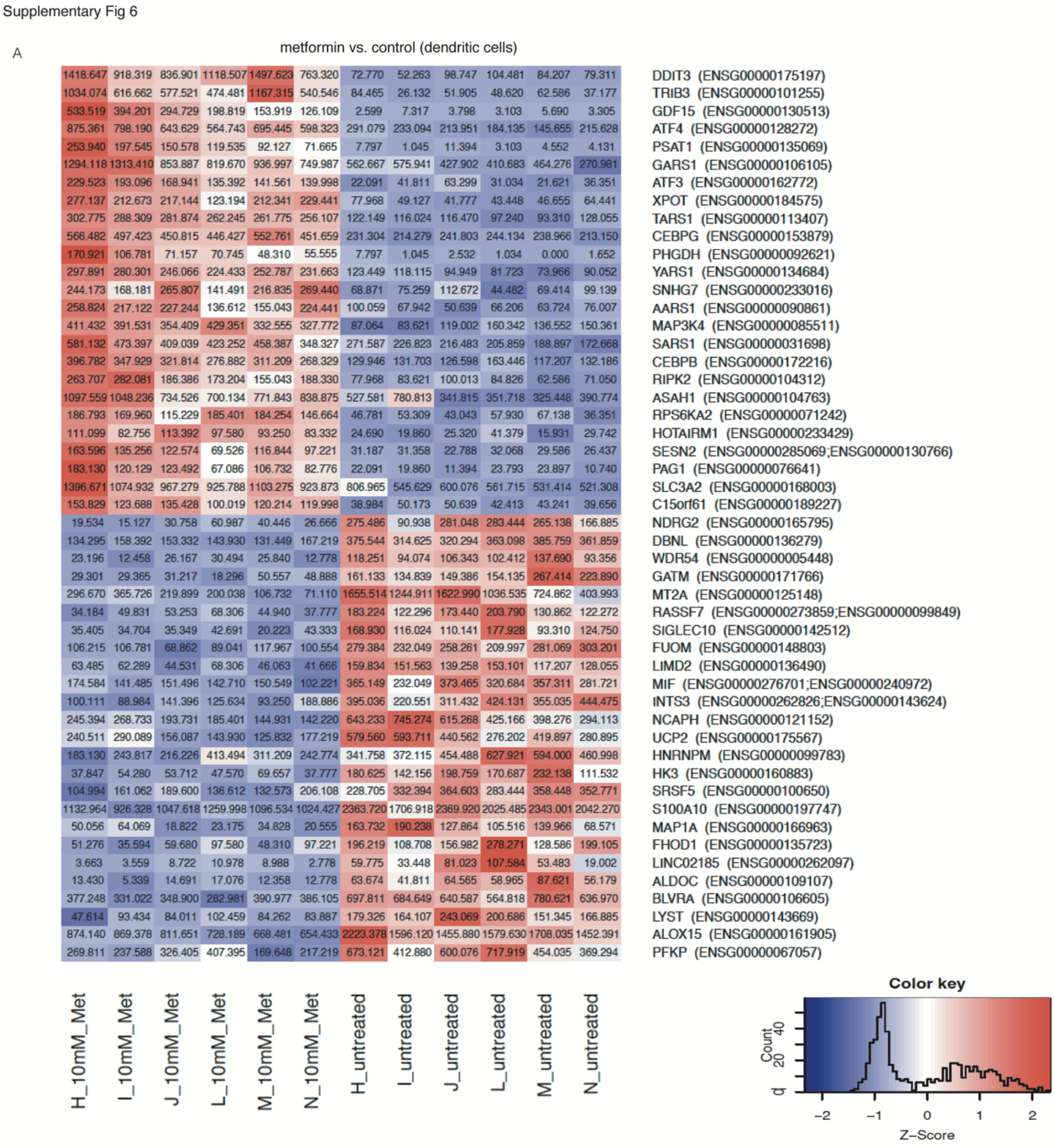


**Supplementary Fig 6. Gene expression chances in PBMC-derived DCs following metformin treatment a,** 25 upregulated (red) and downregulated (blue) differentially expressed genes of PBMC-derived human DCs in response to 24hr treatment of 10mM metformin from bulk RNA sequencing.

Supplementary Fig 7

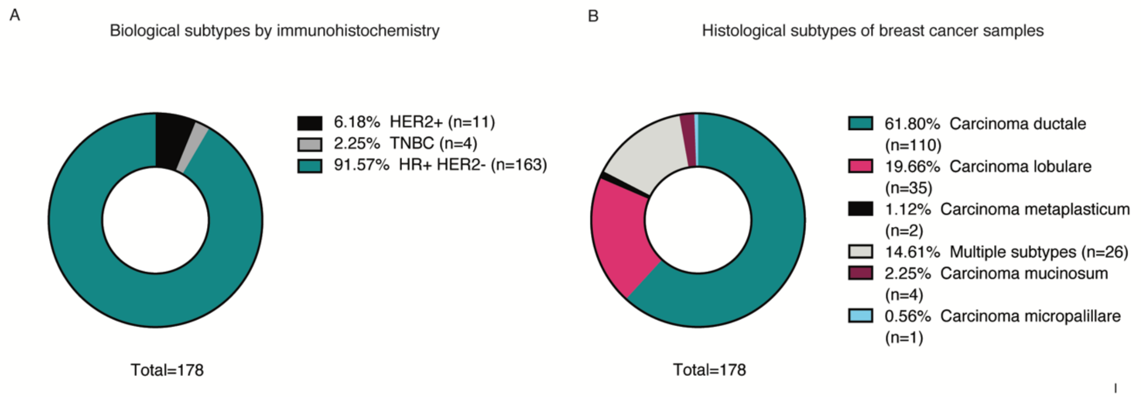


**Supplementary Fig. 7. Breast cancer patient molecular and histological subtypes a,** molecular subtypes of patients from which PDECs were derived for this study. HER2+ = ER-, PR-, HER2+; TNBC = ER-, PR-, HER2-, Luminal = ER+, PR +/-, HER2 +/-. **b,** histological characterization of patients from which PDECs were obtained for this study.

**Supplementary table 1: Flow cytometry antibodies**

| Target | Color | Species | Target | Clone | Catalog | Company |
|---|---|---|---|---|---|---|
| CD103 | unconjugated | Rabbit | | EPR22590-27 | ab224202 | abcam |
| CD11b | AlexaFluor488 | Rat | Human | M1/70 | 557672 | BD |
| CD11c | FITC | | Human | 3.9 | 301604 | BioLegend |
| CD11c | PE | Mouse | Human | 3.9 | 565910 | BD |
| CD11c | unconjugated | | | | PA5-90208 | Invitrogen |
| CD11c | Pe-Cy5 | Mouse | Human | 3.9 | 301610 | Biolegend |
| CD123 | PE-Cy7 | Mouse | Human | 7G3 | 560826 | BD |
| CD123 | BV785 | Mouse | Human | 6H6 | 2130160 | Sony |
| CD14 | BV480 | Mouse | Human | MøP9 | 566141 | BD |
| CD14 | BV510 | Mouse | Human | MøP9 | 563079 | BD |
| CD14 | APC | Mouse | Human | | 555399 | BD |
| CD14 | Pacific blue | Mouse | Human | M5E2 | 301828 | Biolegend |
| CD15 | PerCP-Cy5.5 | Mouse | Human | HI98 | 560828 | BD |
| CD16 | PE | Mouse | Human | B73.1 | 561313 | BD |
| CD163 | BV650 | Mouse | Human | GHI/61 | 563888 | BD |
| CD163 | BV786 | Mouse | Human | GHI/61 | 741003 | BD |
| CD163 | FITC | Mouse | Human | GHI/61 | 563697 | BD |

| | | | | | | |
|---|---|---|---|---|---|---|
| CD163 | BV650 | Mouse | Human | GHI/61 | 563888 | BD |
| CD183 | AlexaFluor488 | Mouse | Human | 1C6/CXCR3 | 558047 | BD |
| CD184 | APC | Mouse | Human | CXCR4 | 560936 | BD |
| CD19 | PE-CF594 | Mouse | Human | HIB19 | 562321 | BD |
| CD19 | BV605 | Mouse | Human | HIB19 | 302244 | Biolegend |
| CD1c | PerCP-Cy5.5 | Mouse | Human | F10/21A3 | 565423 | BD |
| CD1c | APC | | | | | |
| CD206 | APC-Cy7 | | Human | 15-2 | 321120 | BioLegend |
| CD206 | PE | Mouse | Human | | 555954 | BD |
| CD3 | PerCP-Cy5.5 | Mouse | Human | SK7 | 332771 | BD |
| CD3 | Pe-Cy5.5 | | | | | |
| CD4 | PE-Cy7 | Mouse | Human | RPA-T4 | 560649 | BD |
| CD45 | APC-H7 | Mouse | Human | 2D1 | 560178 | BD |
| CD45 | Pacific orange | Mouse | Human | 2D1 | PO-160-T100 | EXBIO |
| CD45 (NCL-L-LCA) | unconjugated | Mouse | Human | | NCL-L-LCA | Novocastra |
| CD56 | BV421 | Mouse | Human | NCAM16.2 | 562751 | BD |
| CD56 | APC-Cy7 | | Human | HCD56 | 318332 | BioLegend |
| CD56 | APC-Cy7 | | | | | |
| CD64 | PE | | Human | 10.1 | 305008 | BioLegend |
| CD64 | PE | | | | | |
| CD66b | AlexaFluor647 | Mouse | Human | G10F5 | 561645 | BD |
| CD69 | APC | | Human | L78 | 340560 | BD |
| CD8 | BV510 | Mouse | Human | SK1 | 563919 | BD |
| CD80 | BV510 | Mouse | Human | L307.4 | 563084 | BD |
| CD80 | AlexaFluor647 | | Human | 2D10 | 305216 | BioLegend |
| CD86 | PE-Cy7 | Mouse | Human | 2331 | 561128 | BD |
| CD86 | PE-Cy7 | | Human | IT2.2 | 305422 | BioLegend |
| CD86 | Pe-Cy7 | Mouse | Human | IT2.2 | 305422 | Biolegend |
| CD8a | unconjugated | Rabbit | | EPR20305 | ab209775 | abcam |
| CD8a | Pe-Cy5 | | Human | HIT8a | 300909 | BioLegend |
| EpCAM | PerCP-Cy5.5 | | Human | EBA-1 | 347199 | BD |
| Fc epsilon R1 alpha | FITC | Mouse | Human | CRA1 | 130-117-361 | Miltenyi |
| FOXP3 | PE | Mouse | Human | 259D/C7 | 560046 | BD |
| GZMB | PE-Cy7 | Mouse | Human | GB11 | 561142 | BD |
| HLA DR | AlexaFluor 594 | Mouse | Human | L243 | NB100-77855AF594 | Novus Biologicals |
| HLA-DR | PE | | Human | L243 | 307606 | BioLegend |
| HLA-DR | APC | Mouse | Human | | 559866 | BD |

| | | | | | | |
|---|---|---|---|---|---|---|
| HLA-DR | APC-R700 | Mouse | Human | G46-6 | 565127 | BD |
| LAG-3 | PE | | Human | 3DS223H | 12-2239-42 | eBioscience |
| Mannose Receptor | AlexaFluor 700 | Mouse | Human | 15 2 | 321132 | Biolegend |
| PD1 | FITC | Mouse | Human | MIH4 | 557860 | BD |
| PDL1 | PE-Cy7 | Mouse | Human | MIH1 | 558017 | BD |
| PDL1 | PerCP-Cy5.5 | Rat | Human | MIH5 | NBP1-43262PECY55 | NovusBio |
| PDL2 (CD273) | APC | Mouse | Human | MIH18 | 557926 | BD |

### Supplementary table 2: IHC antibodies

| Target | Fluorochrome | Species | Target | Dilution | Catalog | Company |
|---|---|---|---|---|---|---|
| FoxP3 | TSA-488 | Mouse | Human | 1:200 | ab20034 | Abcam |
| CD3 | TSA-555 | Rabbit | Human | 1:750 | MA5-14482 | Invitrogen |
| CD8 | Alexa-647 | Mouse | Human | 1:300 | M7103 | DAKO |
| CD4 | Alexa-750 | Rabbit | Human | 1:25 | ab133616 | Abcam |
| CD45 | Alexa-647 | Rabbit | Human | 1:100 | CST13917 | Cell signaling technology |
| PanEpi Cocktail (PanCK, E-cadherin) | Alexa-750 | Mouse | Human | 1:150<br>1:100<br>1:200 | ab7753,<br>MA5-13156,<br>610182 | Abcam<br>Invitrogen<br>BD |
| CD8a | Unconjugated | Rabbit | Mouse | 1:2000 | ab209775 | Abcam |
| CD103 | Unconjugated | Rabbit | Human/ Mouse | 1:5000 | ab224202 | Abcam |
| c-MYC | Unconjugated | Rabbit | Human/ Mouse | 1:200 | ab32072 | Abcam |

### Supplementary table 3: Primers

| Name | Primer nucleotide sequence (5'-3'): |
|---|---|
| Human Arg1 Forward | GAAAGGCTGGTCTGCTTGAG |
| Human Arg1 Reverse | CACAGACCTTGGATTCTTCACA |
| Human iNOS Forward | AATCTCTGGTCAAGCTGGATG |
| Human iNOS Reverse | GCAAGATTTGGACCTGCAAG |
| Human Lamp3 Forward | TTGACCGTCTCAGATCCAGA |
| Human Lamp3 Reverse | CTCTGTTCACTCACGCACTT |
| Human CIITA Forward | TACTCAGAACCCGACACAGA |
| Human CIITA Reverse | CCGATCACTTCATCTGGTCC |
| Human IFNy Forward | TTTAATGCAGGTCATTCAGATG |
| Human IFNy Reverse | AGACAATTTGGCTCTGCATT |
| Human CD47 Forward | TAGATCCGGTGGTATGGATG |
| Human CD47 Reverse | ATATTCACCTGGGACGAAAG |
| Human PRF1 Forward | CGCCTACCTCAGGCTTATCTC |
| Human PRF1 Reverse | CCTCGACAGTCAGGCAGTC |
| Human GZMB Forward | CCCTGGGAAAACACTCACACA |
| Human GZMB Reverse | CACAACTCAATGGTACTGTCGT |
| Human GAPDH Forward | CTCTGCTCCTCCTGTTCGAC |
| Human GAPDH Reverse | GCCCAATACGACCAAATCC |

| | |
|---|---|
| Human ActB Forward | CTTCACCACCACGGC |
| Human ActB Reverse | CCATCTCTTGCTCGAAG |
| Human PUM1 Forward | GCCCCAGTCTTTGCAATTTA |
| Human PUM1 Reverse | AATCACTCGGCAGCCATAAG |
